# Diet-microbiome interactions promote enteric nervous system resilience following spinal cord injury

**DOI:** 10.1101/2024.06.06.597793

**Authors:** Adam M. Hamilton, Lisa Blackmer-Raynolds, Yaqing Li, Sean Kelly, Nardos Kebede, Anna Williams, Jianjun Chang, Sandra M. Garraway, Shanthi Srinivasan, Timothy R. Sampson

## Abstract

Spinal cord injury (SCI) results in a plethora of physiological dysfunctions across all body systems, including intestinal dysmotility and atrophy of the enteric nervous system (ENS). Typically, the ENS has capacity to recover from perturbation, so it is unclear why intestinal pathophysiologies persist after traumatic spinal injury. With emerging evidence demonstrating SCI-induced alterations to the gut microbiome composition, we hypothesized that modulation of the gut microbiome could contribute to enteric nervous system recovery after injury. Here, we show that intervention with the dietary fiber, inulin prevents ENS atrophy and limits SCI-induced intestinal dysmotility in mice. However, SCI-associated microbiomes and exposure to specific SCI-sensitive gut microbes are not sufficient to modulate injury-induced intestinal dysmotility. Intervention with microbially-derived short-chain fatty acid (SCFA) metabolites prevents ENS dysfunctions and phenocopies inulin treatment in injured mice, implicating these microbiome metabolites in protection of the ENS. Notably, inulin-mediated resilience is dependent on signaling by the cytokine IL-10, highlighting a critical diet-microbiome-immune axis that promotes ENS resilience following SCI. Overall, we demonstrate that diet and microbially-derived signals distinctly impact recovery of the ENS after traumatic spinal injury. This protective diet-microbiome-immune axis may represent a foundation to uncover etiological mechanisms and future therapeutics for SCI-induced neurogenic bowel.

## Introduction

Spinal cord injury (SCI) disrupts numerous aspects of physiology, most notably impacting autonomic and voluntary systems at or below the level of lesion^1, 2^. The gastrointestinal (GI) tract is no exception. Up to 60% of injured persons present with myriad intestinal symptoms including constipation, fecal incontinence, and slowed GI transit time, collectively termed neurogenic bowel dysfunction (NBD)^3, 4^. NBD consistently ranks among the most burdensome aspects of SCI – often prioritized over lower limb function^5, 6^; with complications resulting in significant hospitalizations and mortalities^7, 8^. Rodent injuries recapitulate SCI-induced enteric dysfunctions^9, 10^, including reductions in GI transit time, and enteric nervous system (ENS) atrophy and hypoactivity^11–17^. Despite the ability of the healthy ENS to recover following perturbation^18^, these pathologies persist^19, 20^. Thus, we hypothesize that the intestinal environment that arises after injury may inhibit ENS recovery.

One component of the intestinal environment is the indigenous gut microbiome. This complex microbial community plays pivotal roles in host physiology, including the development and modulation of the ENS^21–23^. Emerging data highlight an association between the gut microbiome and locomotor recovery or pain after SCI^24–26^. However, the potential pathogenicity of the injury-associated microbiome and precise contributions to recovery are unknown. After SCI, the gut microbiome composition is markedly shifted^27^. While there is not yet a characteristic gut microbiome composition across SCI studies, some generalities are emerging. A reduction of short chain fatty acid (SCFA)-producing bacteria appears in both experimental and human injuries^27^. SCFAs, produced by bacterial fermentation of dietary fiber, modulate the host intestinal environment, including promoting beneficial immune responses and GI motility^28^. Despite the microbiome’s ability to promote enteric neurogenesis in healthy individuals^21, 22^, it is unknown whether the post-injury gut microbiome contributes to recovery of the GI tract following SCI. Dietary fiber interventions enhance the production of SCFAs and promote microbiome resilience which impact ENS physiology^29, 30^. We therefore sought to determine whether the microbiome-fermented dietary fiber inulin can limit enteric pathologies following SCI.

Here, we used a murine T9/10 contusion model of SCI, that recapitulates hallmark aspects of NBD^31^. We find that inulin supplementation improves gut transit and prevents ENS atrophy. Employing gnotobiotic approaches, we observe that neither the injury-associated microbiome itself nor probiotic interventions with bacterial species lost after SCI were sufficient to modulate enteric pathologies. However, we identify the microbially-derived SCFA metabolite, butyrate, as one factor that limits SCI-induced enteric pathology, suggesting diet-microbe interactions facilitate ENS recovery. Since both inulin and SCFA interventions specifically increased the production of the multi-trophic cytokine IL-10, we investigated its involvement in enteric resilience post-injury. Indeed, injured mice lacking the IL-10 receptor did not respond to dietary inulin, highlighting this immune pathway in prevention of ENS atrophy. Overall, our data elucidate a critical microbiome-neuroimmune interaction elicited by diet that facilitates enteric neuronal and physiological resilience, prevents intestinal dysmotility, and improves enteric pathologies post-SCI.

## Results

### Inulin limits NBD pathology following mid-thoracic SCI in mice

As observed in human injuries, thoracic SCI in rodents triggers significant enteric dysfunction, including colonic dysmotility, ENS atrophy, and microbiome alterations^11–17^. To investigate diet and microbiome dependent effects on SCI-induced NBD, we performed severe contusion injuries (∼70 kilodyne, ∼1.5mm displacement) at the T9/T10 mid-thoracic spinal cord (vertebrae T7-T8)^32^ in mice, in comparison to identical sham laminectomies. Injured animals were immediately provided with either water (vehicle) or water containing 1% soluble inulin (Fig. 1A)- a fiber, dose, and route previously demonstrated to promote resilience of the microbiome^33^. Regardless of dietary intervention, all injured mice quickly displayed significant hind-limb locomotor dysfunction, as measured by the Basso Mouse Scale (BMS)^34^ and weight loss (Fig. 1B, C). At 14-days post-injury (dpi), a timepoint of early recovery rather than acute injury, injured mice displayed slowed intestinal motility, which was significantly improved in those treated with dietary inulin (Fig. 1D). Total intestinal transit is regulated by signals extrinsic to the GI tract (spinal and hormonal, for instance) as well as intrinsically by the ENS. To test the functionality of the ENS in the colon, as the site where inulin would largely be fermented, we recorded colonic contractions in an *ex vivo* preparation. We found that colons derived from injured mice treated with inulin displayed a higher frequency of contractions in the distal colon (Fig. 1E-G). Thus, we hypothesized that inulin acts on intrinsic intestinal physiologies to prevent SCI-induced dysmotility.

**Figure 1.**
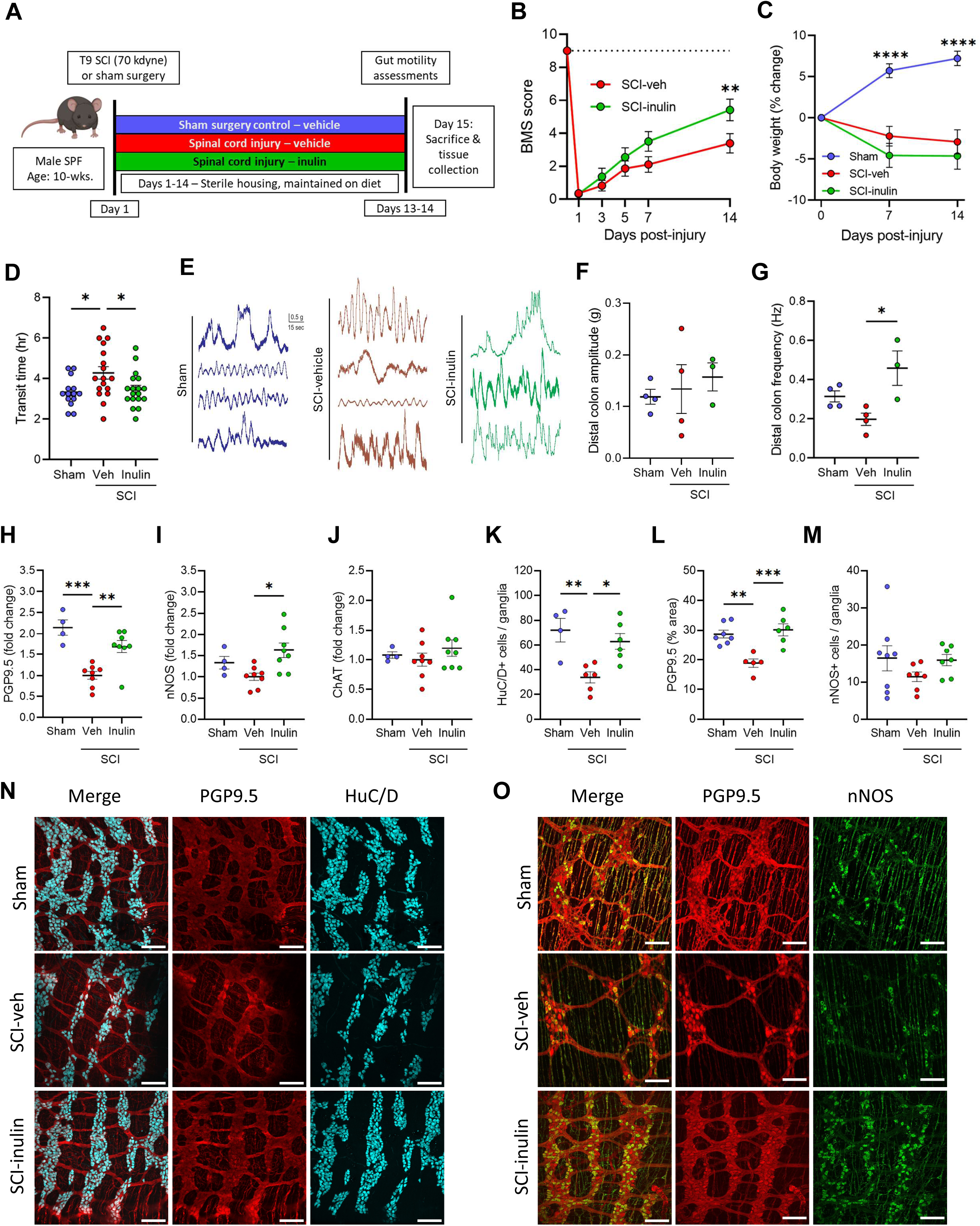
Dietary inulin rescues SCI-induced enteric neuropathy and neurogenic bowel. **A**, Experimental overview: male wild-type mice receive T9 laminectomy (Sham) or laminectomy with 70kilodyne contusive spinal cord injury (SCI), with either a standard diet (Veh) or inulin (Inulin). **B,** Hindlimb locomotor score, as assessed by Basso mouse scale (BMS), with dashed line at 9 representing no deficits, indicative or sham or uninjured locomotor function. **C,** Percent changes in body weight relative to pre-injury weight. **D,** Quantification of total intestinal transit time. **E**, Representative traces of *ex vivo* colonic contractility. **F-G**, Average amplitude (**F**) and frequency (**G**) recorded from distal colon. **H-J**, Colonic protein levels measured via western blot for protein gene product 9.5 (PGP9.5) (**H**), neuronal nitric oxide synthase (nNOS) (**I**), and choline acetyltransferase (ChAT) (**J**). **K-M**, IF quantification showing the number of HuC/D^+^ cells per ganglia (**K**), percentage of PGP9.5^+^ area (**L**), and nNOS^+^ cells per ganglia (**M**). **N, O,** Representative images of myenteric plexus where cyan=HuC/D, red=PGP9.5, and green=nNOS. Asterisks in (**C**) indicate differences between sham and both SCI groups. Each circle represents individuals excluding (**B, C**), where each circle is the mean within that experimental group. IF data points are the averaged values of 2-7 images per mouse with N=4-8 mice per group. N=11-14 (**B**), N=18-19 (**C**), N=14-18 (**D**), N=3-4 (**F, G**), N=4-8 (**H-J**), N=4-8 (**K-M**). * *P* < 0.05, ** *P* < 0.01, *** *P* < 0.001, **** *P* < 0.0001. Data are shown as mean ± SEM and compared by repeated measures 2-way ANOVA with post-hoc Tukey’s test (for **B, C**) and by one-way ANOVA with post-hoc Tukey’s test (for **D, F-M**). Dashed line in (**B**) indicates maximum possible score for BMS of 9.

Subsequent molecular characterization of enteric neuronal markers in colonic tissues post-SCI validated prior reports of ENS dysfunction and colonic remodeling ^11–17, 35^. We observed a substantial decrease in PGP9.5 protein post-SCI, as well as a qualitative decrease in nNOS, but not in ChAT by western blot analysis (Fig. 1H-J). Dietary inulin intervention resulted in substantial increases of PGP9.5 and nNOS protein levels (Fig. 1H, I), indicating it limits the loss of these markers and highlighting the sensitivity of nitrergic enteric signaling in the post-injury environment. Correspondingly, immunofluorescence imaging of the colonic myenteric plexus revealed a substantial atrophy of enteric neurons post-SCI (Fig. 1K-O). This includes both a loss of HuC/D^+^ cells within the myenteric ganglia and a diminished total area of ganglia and neuronal processes (PGP9.5), but no significant difference in nNOS^+^ cells. This neuronal loss and ganglia atrophy was substantially limited in injured mice receiving inulin (Fig. 1K-O). Thus, this dietary fiber intervention maintains total intestinal transit following SCI, corresponding to a protection from myenteric neuronal loss.

### Dietary inulin prevents SCI-triggered gut dysbiosis

The gut microbiome readily responds to dietary inputs, and dietary fiber specifically promotes its resiliency^33^. SCI in rodents has demonstrated progressive changes to overall gut microbiome composition beginning at ∼14- dpi^24^. We therefore investigated whether inulin intervention prevented SCI-induced microbiome compositional alterations using 16S rRNA profiling. At 14-dpi, we note limited impacts to alpha-diversity of the fecal microbiome (Fig. 2A, B), but a substantial alteration to beta diversity within the community (Fig. 2C, D and Supplementary Tables 1, 2). A fully independent cohort was impacted similarly, although not identically, following SCI (Supplementary Fig. 1 and Supplementary Tables 3-5). Across these independent studies, a small number of bacterial taxa, including *Bacteroides*, *Clostridium*, *Lactobacillus*, and *Turicibacter* were consistently sensitive to SCI, with increased or decreased relative abundances (Fig. 2E and Supplementary Table 3). Inulin intervention did not completely prevent an SCI-induced impact on the microbiome (Fig. 2A-E and Supplementary Fig. 1), and in fact largely created a community structure more unique to itself, rather than sham or vehicle-treated mice. We did observe a select set of SCI-sensitive taxa that were less affected by SCI, when in the presence of inulin. *Bacteroides sp.* and *Lactobacillus johnsonii* were increased by inulin post-SCI, while others such as *Clostridium celatum* and *Turicibacter sanguinis* remained similar to vehicle-treated, injured mice (Fig. 2E-K). These data demonstrate an effect of inulin on post-injury microbiome composition, corresponding to its ability to prevent SCI-triggered behavioral and molecular pathologies in the colon.

**Figure 2.**
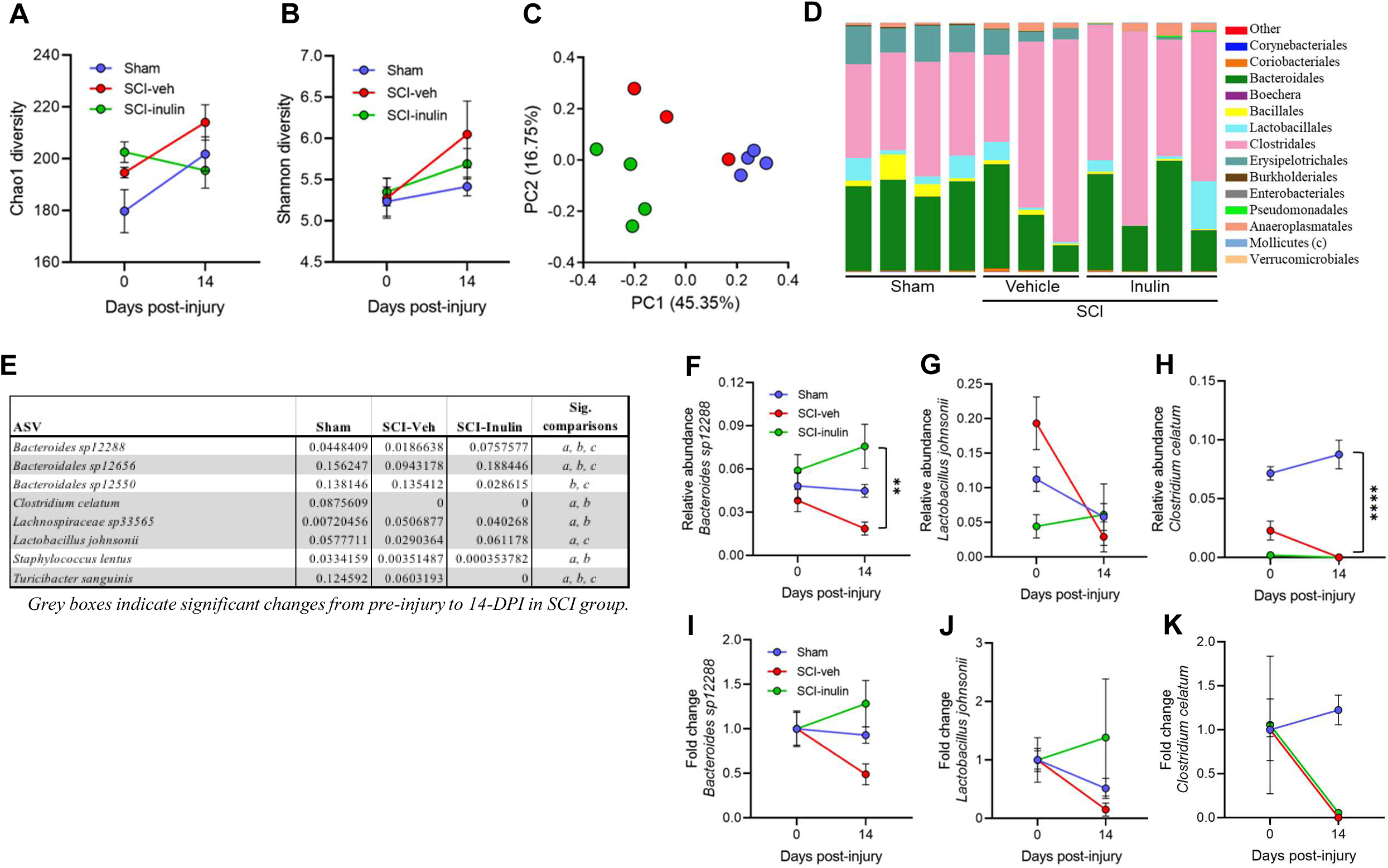
SCI-triggered dysbiosis is prevented by inulin. **A, B,** Pre-surgery to 14-dpi changes in fecal microbiome in male mice. Chao1(**A**) and Shannon (**B**) alpha diversity measures. **C**, Bray-Curtis principal component analysis (PCA) plot. **D**, Order level microbiome composition plot at 14-dpi. **E**, Table highlighting the mean relative abundance of bacterial taxa that differ between experimental groups at 1% FDR, where grey boxes indicate significant changes from pre-injury to 14-dpi in SCI-veh group, and “a, b, & c” indicate significant differences between groups at endpoint. **F-K**, Progressive changes in select taxa represented as relative abundance (**F-H**) and as proportion of pre-injury abundance (**I-K**) for *Bacteroides sp12288* (**F, I**), *Lactobacillus johnsonii* (**G, J**), and *Clostridium celatum* (**H, K**). Asterisks in (**F**) indicate significant difference between SCI-veh and SCI-inulin groups at 14-dpi. Asterisks in (**H**) indicate significant difference between sham and both SCI groups. Each circle represents within group mean, excluding **C**, where each circle represents individuals. N=3-4. ** *P* < 0.01, **** *P* < 0.0001. Data are shown as mean ± SEM and compared by repeated measures 2-way ANOVA with post-hoc Sidak’s test (**A, B**) or Tukey’s test (**F-K**) and by 1% FDR (for **E**).

### Injury- and diet-associated gut microbiomes are not sufficient to induce or prevent NBD

Enteric physiology and neurogenesis are modulated by metabolic and immune responses derived from signals of the gut microbiome^21–23^. Given our finding that dietary fiber could prevent aspects of NBD, we sought to determine if the injury- or diet-associated microbiomes alone were sufficient to impact intestinal transit. We used an injury-naïve, gnotobiotic mouse model, reconstituting fecal microbiomes derived from mice with SCI- with or without inulin- or sham injured controls (Fig. 3A). Surprisingly, there was no recapitulation of donor phenotypes in the recipients’ intestinal transit (Fig. 3B), indicating that the microbes themselves are not sufficient to drive intestinal transit in injury-naïve mice, nor is the benefit of inulin solely due to restructuring of the microbiome composition. Although the fecal microbiome was not sufficient to recapitulate the intestinal motility phenotypes of their donors in recipient mice, a number of colonic and circulating metabolic and inflammatory signals were directly affected by the microbiome (Supplementary Fig. S2A-G). This included elevated levels of colonic CXCL1 and IL-12p70 (Supplementary Fig. S2A-C), as well as reduced levels of c-peptide, TNF, and ghrelin in circulation (Supplementary Fig. S2D-G) in recipients of SCI-derived microbiomes. Abundances of many, but not all, of these circulating factors mirror those found in the microbiome donor animals, including reduced levels of c-peptide and TNF in mice with SCI (Supplementary Fig. S2H-M), providing evidence for microbiome-dependent contributions to these aspects of metabolic and immune function. The microbiome that arises post-SCI therefore is sufficient to differentially modulate physiologic pathways which may affect other pathophysiologies post-injury, despite not being sufficient to modulate intestinal transit in injury-naïve, germ-free mice.

**Figure 3.**
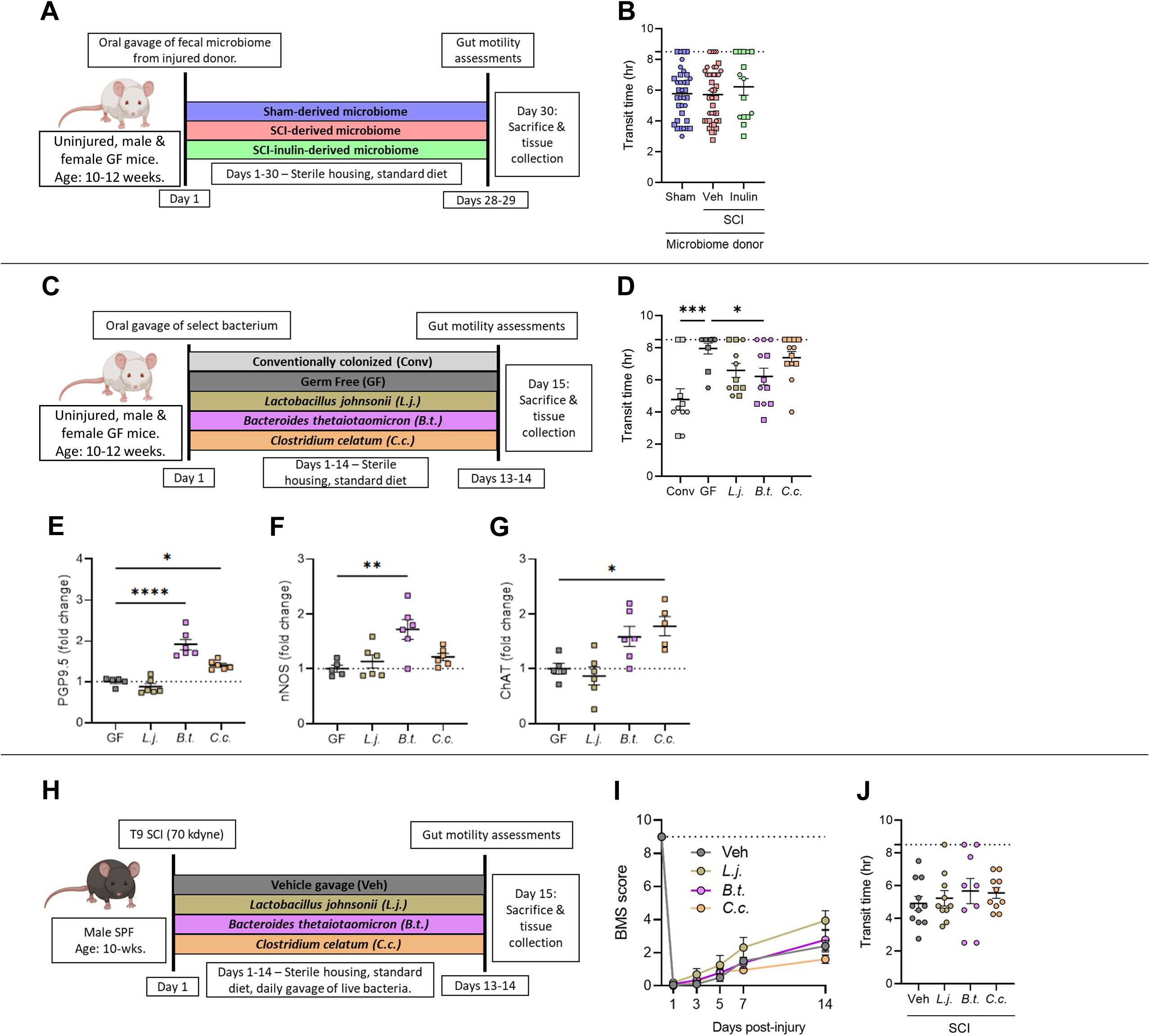
Post-injury and diet-induced microbes differentially impact the ENS. **A**, Experimental overview: male and female, uninjured, wild-type, germ-free mice received fecal microbiomes of donors with laminectomy (sham), SCI (veh) or SCI with inulin (inulin). **B,** Quantification of total intestinal transit time. **C**, Experimental overview: male and female, uninjured, wild-type, germ-free mice are mono-colonized with *Lactobacillus johnsonii* (*L.j.*), *Bacteroides thetaiotaomicron* (*B.t.*), or *Clostridium celatum* (*C.c.*), alongside conventionally colonized (Conv) and germ-free (GF) controls. **D**, intestinal transit time at experimental endpoint. **E-G**, Quantification of proximal colon, from male mice, by western blots for protein gene product 9.5 (PGP9.5) (**E**), neuronal nitric oxide synthase (nNOS) (**F**), and choline acetyltransferase (ChAT) (**G**). **H**, Experimental overview: male, wild-type mice received laminectomy with 70kilodyne contusive spinal cord injury (SCI), followed by daily oral gavage of *L.j., B.t., C.c.*, or vehicle (veh). I, Progressive Basso mouse scale (BMS) scores of mice provided daily probiotic gavage of veh, *L.j., B.t., or C.c.,* with dashed line at 9 representing uninjured locomotor function. **J**, Endpoint intestinal transit time. Each point represents individuals for all but (**I**), where each circle is the average of all mice within that experimental group. (**A-D**) Squares represent males, hexagons represent females. N=15-41 (**B**), N=10-12 (**D**), N=5-6 (**E-G**), N=9-11 (**I-J**). * *P* < 0.05, ** *P* < 0.01, *** *P* < 0.001, **** *P* < 0.0001. Data are shown as mean ± SEM and compared by ordinary one-way ANOVA with post-hoc Tukey’s (**B**) or Dunnett’s (**D-G, J**) tests. Dashed line in (**I**) indicates maximum possible score for BMS of 9.

Since we observe that the early SCI-associated microbiome itself does not trigger intestinal dysmotility in injury-naïve gnotobiotic mice, the loss of certain beneficial organisms may instead result in increased susceptibility to slowed intestinal transit post-injury. We therefore sought to determine whether those microbes decreased following injury could prevent SCI-induced NBD. We tested a panel of organisms consistently depleted following SCI (Fig. 2), in a mono-colonized gnotobiotic setting to establish their sufficiency to modulate intestinal physiological parameters central to SCI-induced NBD (Fig. 3C). We found that mono-colonization of germ-free mice with *Bacteroides thetaiotaomicron* (closely related to *Bacteroides sp.12288*) significantly improved total intestinal transit time compared to the germ-free controls (with impaired intestinal transit due to an underdeveloped ENS^21, 36^) (Fig. 3D), but did not fully recapitulate the transit time of conventionally-colonized mice. Mono-colonization with *Lactobacillus johnsonii* or *Clostridium celatum* showed no significant improvement (Fig. 3D). Notably, western blot analysis of colonic tissue revealed that *B. thetaiotaomicron* substantially increased those ENS markers also impacted by inulin supplementation, PGP9.5 and nNOS, compared to germ-free controls (Fig. 3E-G). *C. celatum* increased both ChAT and PGP9.5, while *L. johnsonii* had no impact on any tested marker (Fig. 3E-G). Thus, some organisms depleted after SCI are sufficient to modulate the production of ENS markers involved in intestinal motility.

Given our observation that *B. thetaiotaomicron* was sufficient to both improve intestinal transit and increase molecular markers of the ENS, including those lost following SCI, we next set out to test whether this organism could differentially prevent the onset of NBD following SCI, similar to the action of dietary inulin. Following SCI, mice were orally gavaged with one of the three select taxa, *B. thetaiotaomicron, L. johnsonii*, or *C. celatum* (Fig. 3H), which each displayed differential ENS outcomes in the mono-colonized system (Fig. 3C-G). While other probiotic species (specifically a cocktail of *Lactobacillus sp.* and *Bifidobacterium sp.*) have been observed to improve locomotor recovery in a similar SCI model^24^, supplementation with any of these three organisms, that we noted to be specifically depleted following SCI, did not result in improved locomotor outcomes (Fig. 3I). Despite its ability to modulate intestinal physiology in a mono-colonized paradigm, *B. thetaiotaomicron* exposure did not improve transit time in injured mice compared to vehicle-treated controls (Fig. 3J). In addition, neither *L. johnsonii* nor *C. celatum* exposure modulated intestinal transit in mice with SCI (Fig. 3J). Overall, these data suggest that while some members of the gut microbiome are sufficient to modulate aspects of the ENS and intestinal motility in a mono-colonized and injury-naïve setting, SCI- and inulin-associated microbial communities are not themselves sufficient to recapitulate, or reduce, NBD in the absence of injury. Furthermore, gut exposure to individual taxa that are preferentially depleted post-SCI did not prevent intestinal dysmotility, suggesting a diet-microbiome interaction in the facilitation of inulin’s beneficial effects.

### IL-10 signaling is necessary to limit intestinal dysmotility post-SCI

Certain cytokines can promote neurogenesis and limit neurodegenerative processes^37, 38^. We therefore comprehensively assessed a panel of intestinal cytokines post-SCI via multi-plex ELISA. At 14-dpi, we did not observe any significant differences in colonic cytokine levels between sham and injured mice treated with vehicle (Supplementary Fig. S3A). Inulin intervention in mice with SCI, however, resulted in a substantial increase in the pleiotropic cytokine IL-10 in the colon relative to vehicle-treated, SCI mice (Fig. 4A), as well as decreased levels of the pro-inflammatory cytokine IL-1β (Supplementary Fig. S3B), with limited diet-induced effects on other cytokines post-SCI (Supplementary Fig. S3A). We therefore hypothesized that IL-10 signaling may be a necessary component to mediate inulin-induced enteric resilience following SCI.

**Figure 4.**
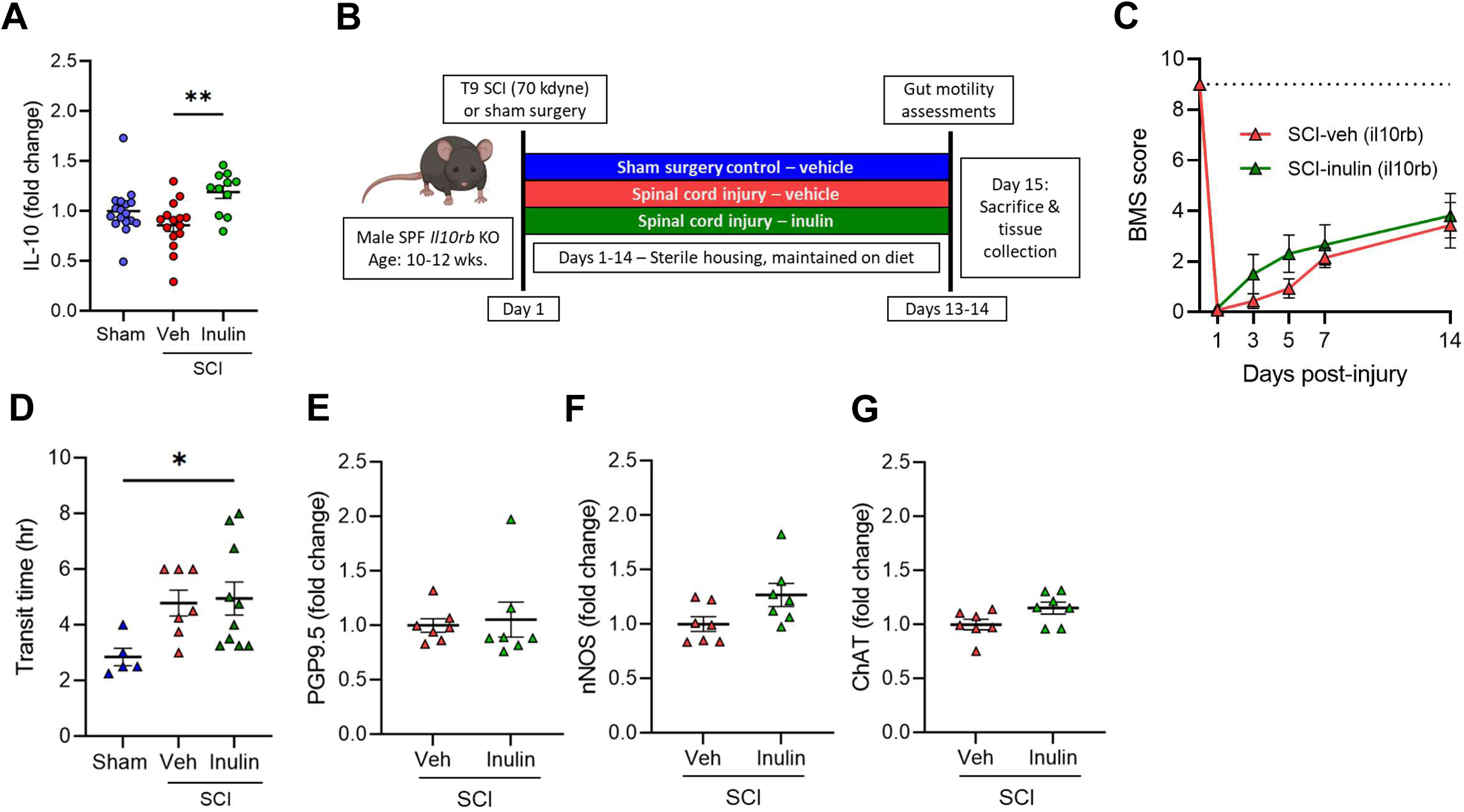
IL-10 signaling is necessary for inulin-mediated resilience to NBD. **A**, Concentration of colonic IL-10 by ELISA from male mice with laminectomy (sham), SCI (veh), or SCI with inulin-supplemented diet (inulin). **B**, Experimental overview: male *IL10rb* KO mice undergo laminectomy at T9 (Sham) or laminectomy with 70kilodyne contusive spinal cord injury (SCI) and receive either standard diet (Veh) or inulin (Inulin). **C**, Progressive Basso mouse scale (BMS) scores with dashed line at 9 representing uninjured locomotor function. **D**, intestinal transit time at endpoint. **E-G**, Quantification of colonic protein gene product 9.5 (PGP9.5) (**E**), neuronal nitric oxide synthase (nNOS) (**F**), and choline acetyltransferase (ChAT) (**G**) by western blot. Each point represents individuals for all but (**C**), where each circle is the average of all mice within that experimental group. N=11-17 (**A**), N= 5-10 (**C-G**) * *P* < 0.05, ** *P* < 0.01. Data are shown as mean ± SEM and compared by ordinary one-way ANOVA with post-hoc Tukey’s (**A, D**), 2-way ANOVA (**C**) or two-tailed unpaired t-test (**E-G**). Dashed line in (**C**) indicates maximum possible score for BMS of 9.

We thus directly tested the contribution of IL-10 signaling to resilience post-SCI using IL10rb KO mice, which lack a key subunit in the IL-10 receptor (Fig. 4B). Unlike WT mice (Fig. 1), inulin supplementation did not improve any post-SCI locomotor or enteric outcomes in IL10rb KO mice (Fig. 4C-G). Inulin intervention could not limit SCI-induced deficits in intestinal transit with both diet and vehicle groups displaying impaired transit post-injury (Fig. 4D). In addition, colonic PGP9.5 remained unchanged despite inulin supplementation in IL10rb KO mice post-SCI (Fig. 4E), with subtle alterations to other colonic ENS markers following injury (Fig. 4F, G). Thus, the IL-10 signaling pathway is central to both intestinal behaviors and molecular resilience to SCI-induced enteric pathologies and is activated by inulin intervention.

### SCFA signaling is sufficient to improve SCI-induced dysmotility

Inulin is fermented by the gut microbiome to SCFAs^39^, which have broad roles in host physiology, including the ability to promote a beneficial anti-inflammatory intestinal environment^30^. We observed that the injury-associated microbes themselves were not solely responsible for either the pathogenesis of SCI-induced NBD nor its inulin-mediated prevention (Fig. 3). We therefore sought to determine whether the microbiome-derived SCFA metabolites of inulin were mediating the limitation of SCI-triggered intestinal dysmotility. Fecal SCFA abundances between treatment groups were indeed altered in a diet-dependent manner (Fig. 5A-C). Concurrently, we also noted a significant increase in the SCFA receptor FFAR2 in colonic tissue of inulin-treated mice with SCI, indicating an increased physiological response to SCFAs (Fig. 5D). This colonic FFAR2 response appeared specific, since we observed no alteration to other SCFA receptors, including FFAR3 and GPR109 (Fig. 5E, F). Similar effects on fecal SCFA abundances and colonic FFAR2 production following inulin intervention in IL10rb KO mice were also observed (Supplementary Fig. S3C-F) We therefore hypothesized that microbiome-derived SCFA signaling is involved in limiting intestinal pathologies post-SCI, rather than the microbiome composition *per se*, and that these are upstream of IL-10 signaling pathways.

**Figure 5.**
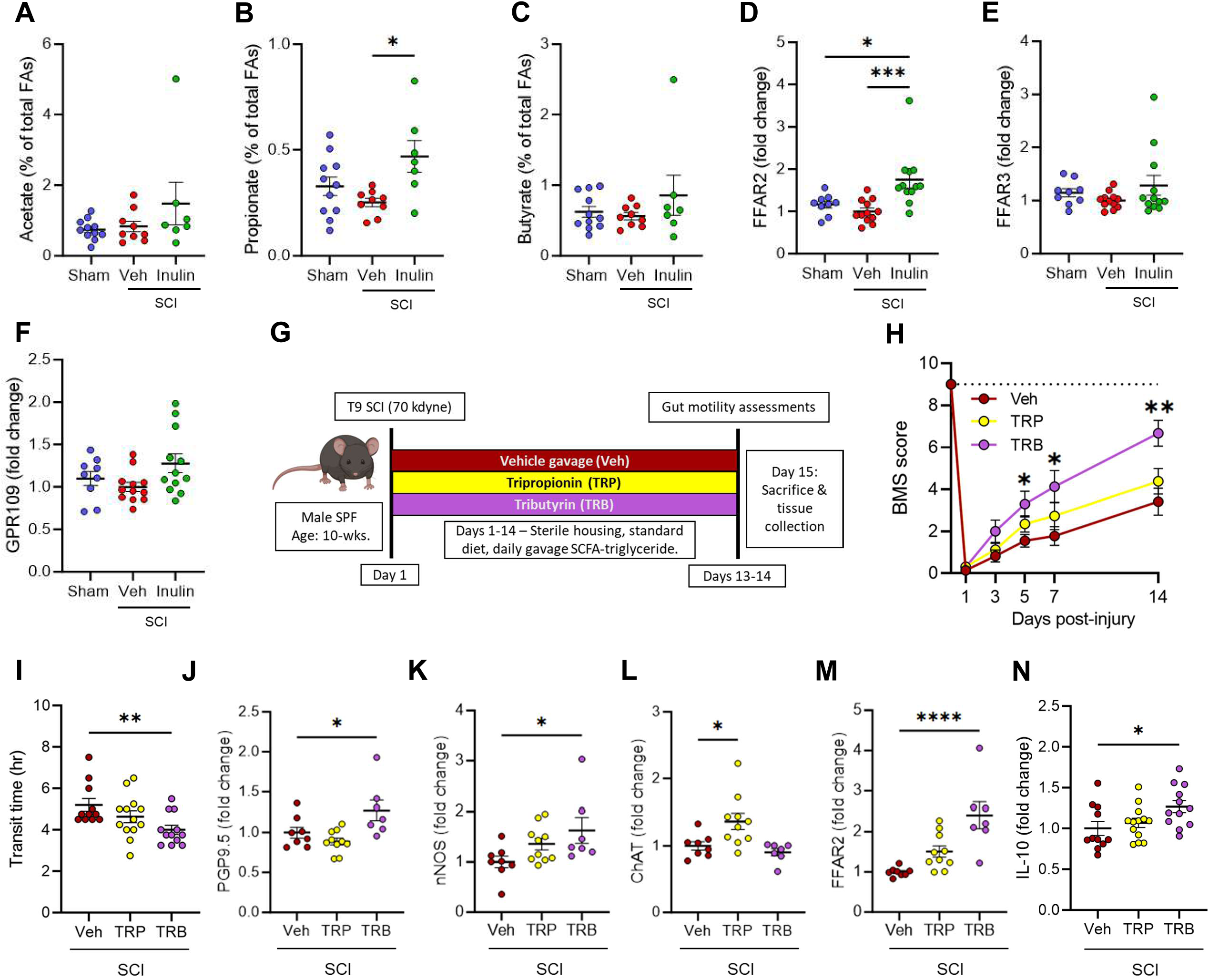
Intestinal SCFA signaling is impacted post-SCI and promotes rescue of NBD. **A-C,** Quantification of indicated short-chain fatty-acids at endpoint fecal pellets from male mice with sham laminectomy (sham), or with SCI (SCI-veh) or inulin-supplemented diet (SCI-inulin). **D-F**, Quantification of colonic free fatty acid receptor 2 (FFAR2) (**D**), free fatty acid receptor 3 (FFAR3) (**E**), and G-protein-coupled receptor 109 (GPR109) (**F**) by western blot. **G**, Experimental overview: male wild-type mice receive laminectomy with 70kilodyne contusive spinal cord injury (SCI) followed by daily gavage of vehicle (veh), tripropionin (TRP), or tributyrin (TRB). **H**, Progressive Basso mouse scale (BMS) scores, with dashed line at 9 uninjured locomotor function. **I**, intestinal transit time at experimental endpoint. **J-M**, Quantification of colonic protein gene product 9.5 (PGP9.5) (**J**), neuronal nitric oxide synthase (nNOS) (**K**), choline acetyltransferase (ChAT) (**L**), and free fatty acid 2 (FFAR2) (**M**) by western blot. **N**, Concentration of serum IL-10 by ELISA. Each point represents individuals for all but (**H**), where each circle is the average of all mice within that experimental group. N=7-11 (**A-C**), N=9-12 (**D-F**), N=11-13 (**H, I, N**), N=7-10 (**J-M**). * *P* < 0.05, ** *P* < 0.01, *** *P* < 0.001, **** *P* < 0.0001. Data are shown as mean ± SEM and compared by ordinary one-way ANOVA with post-hoc Tukey’s (**A-F**) or Dunnett’s (**I-N**) tests, and by 2-way ANOVA with post-hoc Dunnett’s multiple comparison test (**H**). Dashed line in (**H**) indicates maximum possible score for BMS of 9.

To determine a contribution for SCFA signaling in protection against SCI-triggered NBD, we assessed two triglycerides-tripropionin and tributyrin. These molecules are absorbed in the small intestine and metabolized to the SCFAs propionate and butyrate, respectively, by host metabolic processes, rather than the gut microbiome^40^. Immediately following T9/10 contusion SCI, and daily thereafter, injured mice were provided with daily oral gavages containing ∼20mg of tripropionin (TRP) or tributyrin (TRB) diluted in corn oil, in comparison to corn oil vehicle alone (Veh) (Fig. 5G). Tributyrin-treated mice displayed significantly improved locomotor outcomes and intestinal transit, when compared to vehicle-treated mice (Fig. 5H, I). While TRP treatment had little effect on nNOS or PGP9.5 in colonic tissues by western blot analysis, these markers were significantly increased in mice treated with TRB (Fig. 5J, K), phenocopying inulin intervention. Interestingly, TRP, but not TRB, treatment resulted in an increase in colonic ChAT, relative to vehicle controls (Fig. 5L). However, given the concomitant rescue of total intestinal transit time by TRB, we conclude that TRB-derived signaling limits SCI-induced ENS dysfunction and dysmotility, similarly to inulin.

We additionally note an increase in colonic FFAR2 in TRB-treated mice (Fig. 5M), as we also observe after inulin treatment (Fig. 5D), but not significantly in mice treated with TRP post-injury. This corresponds with an increase in circulating IL-6 in TRB-treated mice (Supplementary Fig. S3G, H), along with an increase in circulating IL-10 (Fig. 5N and Supplementary Fig. S3G), further implicating this pathway in protection from SCI-induced intestinal dysmotility and specificity of TRB in these outcomes. Therefore, SCFA signaling is sufficient to limit the effects of SCI on intestinal transit and suggests that the therapeutic activity of inulin is through its fermentation to these microbially-dependent products. These beneficial effects occur despite the rapid metabolism of SCFA-triglycerides, with serum half-lives of only a few hours^41^. Indeed, analysis of serum SCFA concentrations revealed no significant differences in circulating SCFAs 14hrs after mice were administered the SCFA prodrugs (Supplementary Fig. S3I-K). Overall, these data indicate that supplementation with inulin increases the resilience of the ENS to SCI-induced atrophy, through an SCFA-mediated and IL-10-dependent pathway, but largely independent of the microbiome composition as a whole, leading to the prevention of SCI-triggered neurogenic bowel.

## Discussion

The panoply of gastrointestinal dysfunctions that arises following SCI are a significant burden for those with injury and their care partners. Despite the fact that the healthy ENS has the capacity to recover after perturbation, those with injury-induced NBD often show chronic pathophysiologies. In this work, we establish that SCI-triggered enteric pathologies are able to be modified by the intrinsic intestinal environment. Slowed intestinal transit, gut microbial dysbiosis, and enteric neuropathy are each prevented following the addition of the dietary fiber inulin into the intestinal environment. Although SCI modifies the gut microbiome composition, our gnotobiotic and probiotic approaches suggest that the presence or absence of specific SCI- or fiber-associated microbes does not contribute to these intestinal pathologies. We provide evidence that inulin and its microbially-derived SCFAs, through an ability to promote IL-10 signaling, are capable of reducing enteric atrophy and promoting intestinal transit following SCI. Together these data highlight a central diet-microbiome interaction that can be modulated to limit SCI-induced NBD in mice.

Some probiotic treatments (*e.g.* VSL#3/Visbiome), have been demonstrated to promote locomotor recovery following SCI^24^. In our approach, we observe that SCI-sensitive microbes are not sufficient to restore colonic motility nor trigger intestinal dysmotility in gnotobiotic models. We conclude that the composition of the microbiome is therefore not solely responsible for chronic enteric pathologies following SCI. It is possible that injury-associated microbes require an intestinal environment shaped by SCI to impart an effect on the host. For instance, increased intestinal permeability post-SCI^24^ may result in translocation of specific microbial molecules more readily. It may also be that the under-developed ENS of germ-free mice lacks the ability to respond to the microbes we introduced. However, we do observe improvements in intestinal transit in certain mono-colonized conditions, suggesting the capacity of some SCI-associated microbes to restore intestinal transit of germ-free mice. Nonetheless, our data support the notion that diet-microbe interactions or microbe-microbe interactions (*i.e.* microbial metabolism or cross-feeding, *etc.*) have the capacity to restore colonic motility after SCI and should be considered in conjunction with probiotic therapy. It is important to note that our study relied on male animals. Since ∼79% of SCI incidences in the United States occur in males, this is a relevant experimental group^42^. While our tests of microbiome impacts were performed in mice across both sexes, we do not note any observable sex-dependent effects in microbiome-dependent transit in those studies. Future studies, powered to compare male and female responses, would be necessary to understand whether these interventions have similar effects regardless of biological sex.

Emerging evidence indicates that the ENS undergoes homeostatic neurogenesis^43^, and can acutely recover following perturbation, such as infection^44^. Intestinal signaling pathways, including serotonergic signaling from enterochromaffin cells, microbial stimulation of neuronal TLR2, and SCFAs have each been shown independently to modulate enteric neurogenesis^21–23^. Enteric nitrergic neurons appear more sensitive to perturbation than other cells in the ENS^45^, which we also observe following SCI, leading to decreased propulsion of luminal contents. Our data suggest that diet-induced SCFAs regulate resilience of enteric nitrergic neurons by broadly creating an anti-inflammatory/neuroprotective intestinal environment, rich in IL-10. While we do not observe injury-dependent impacts on SCFA signaling pathways, our data provide evidence that modulation of SCFA- and IL-10-dependent processes can promote enteric neuron resilience and limit the effects of SCI on intestinal dysmotility.

Through the study of dietary fiber and the microbiome after SCI, we have identified a previously unexplored signaling pathway able to be readily modulated to prevent injury-associated intestinal dysmotility and ENS atrophy. While intestinal serotonergic signaling has been used to treat SCI-induced NBD^46^, FFAR2 activity and IL-10 treatment have not. IL-10, however, has been shown to have therapeutic value in locomotor recovery during systemic or spinal cord application post-injury^38, 47^. Our observations suggest that inulin and TRB increase local and systemic IL-10 and therefore may prevent not only the onset of enteric pathologies, but also improve systemic outcomes. In line with this, we observed significant locomotor recovery in mice treated with inulin or TRB post-SCI (Fig. 1B and Fig. 5H), similar to prior work with SCFAs themselves^48^. Despite the ease of administration, few studies to date have investigated a defined dietary fiber or its metabolites in persons living with SCI^49, 50^. Our data here provide pre-clinical justification for the continued investigation of diet-microbe interactions as therapeutic targets, to prevent and restore bowel function in those with traumatic spinal cord injury.

## Methods

### Animal husbandry

Wild-type C57Bl6/J, IL10rb^-/-^, and DBA/2J mice were obtained from Jackson Laboratory (#000664, 005027, & 000671). Mice were housed within a central vivarium in sterile microisolator cages on static racks, with autoclaved food (Teklad Autoclavable Diet, Cat 2019S) and water provided *ad libitum* with a 12:12hr light-dark cycle for the duration of the studies. All handling and cage changes occurred under a sterilized biosafety cabinet. Experimental procedures were approved by the Institutional Animal Care and Use Committee of Emory University (Protocols 201700855, 201900030, and 201900145). Germ-free DBA/2NTac mice were obtained from Taconic Biosciences (#DBA2) following embryonic rederivation and bred within the Emory Gnotobiotic Animal Core (EGAC). Prior to colonization mice were assessed for sterility by plating fecal pellets under aerobic and anaerobic conditions on non-selective tryptic soy agar with 5% sheep’s blood (Hardy #A10).

Mice were humanely euthanized via open-drop isoflurane overdose in an induction chamber followed by cardiac puncture and exsanguination. Mice were then perfused with ice-cold sterile phosphate buffered saline (PBS) prior to tissue collection. The entirety of the GI tract was removed, and the colon length measured. Approx. one cm portions of tissue were taken from the proximal colon, flash-frozen in liquid nitrogen, and stored at −80°C until future analysis. The remaining intestinal tissue was fixed in 4% paraformaldehyde (PFA) overnight at 4°C, then transferred to PBS with 0.01% sodium azide for long-term storage.

### Spinal cord injury

Male mice were deeply anesthetized with 3% isoflurane, using oxygen as a carrier gas, and were maintained on 3% isoflurane for the duration of the surgery. Immediately prior to surgery mice received a subcutaneous injection of meloxicam (5 mg/kg). Under sterile conditions, a dorsal laminectomy (vertebrae T7-T8)^32^ was performed to expose the underlying T9/T10 segment of the thoracic spinal cord. Following laminectomy, mice received a 70 kdyne impact onto the dorsal surface of the spinal cord with an Infinite Horizon impactor device (Precision Systems and Instrumentation, Fairfax Station, VA), as performed previously^51^. Displacement was recorded at 1,554±198µm, indicating a severe contusion for each animal^52^. Care was taken to ensure that dorsal roots were not damaged by the laminectomy or impact, and on-target bilateral bruising of the dorsal spinal cord was verified by examination under a dissecting microscope.

Following surgery, the wound was closed using sterile reflex #7 wound clips, and 0.5ml of 0.9% sterile saline was administered subcutaneously. Sham surgery control mice underwent an identical surgical procedure, including laminectomy, but did not receive a contusive SCI. Mice recovered in sterile cages on a heating pad. Subsequent subcutaneous injections of meloxicam (5mg/kg) were administered each day for 2 days following surgery for all sham and injured animals. Experimenters manually expressed the bladders of injured mice twice daily, ∼12 hours apart. Mice were assessed for impairment of locomotor function at one day post-injury (dpi) using the Basso Mouse Scale (BMS)^34^, to ensure effectiveness of the injury. SCI mice were excluded if they recorded a BMS score of 2 or greater 1-dpi. Sham animals were excluded if they recorded a BMS below 9 at 1-dpi. Following surgical procedures animals were transferred to cages containing sterile absorbent bedding (ALPHA-dri®, Shepherd Specialty Papers) to prevent rashes and abrasions. Additionally, portion cups (Dart Solo SCC100S 1 oz. Squat White Paper Portion Cup) containing sterile moistened chow were changed daily to provide all mice with easy access to food and water. For mice receiving inulin intervention, a 1% inulin (Sigma, 12255) weight/volume solution in water was provided immediately following surgical procedures, replacing standard water, *ad lib*. All handling was performed under a sterile biosafety cabinet.

Mice receiving SCFA triglycerides were given a once daily 200µl oral gavage consisting of diluted tripropionin (Thermo, AC275101000) or tributyrin (Sigma, T8626). Liquid triglycerides were diluted 1:10 in corn oil (Veh) with a single 200µl gavage containing 20µl of triglyceride, corresponding to a dose of 21.6mg (415.7mM) of tripropionin or 20.6mg (341.3mM) of tributyrin per gavage. Initial gavage was provided on the day of surgery with subsequent doses provided daily thereafter.

### Total intestinal transit time

Total intestinal transit was performed via carmine red dye elution^53^. Mice were acclimated to an isolated behavior space for 1hr prior to gavage with 100µl of sterile carmine red dye (6% w/v) (Sigma, C1022) dissolved in 0.5% methylcellulose (Sigma, M7027). Following gavage mice remained in their home cages for 2hrs and were then transferred to individual, sterile cages devoid of bedding and food/water. Every 15min, cages were checked for the presence of a red fecal pellet. Immediately upon discovery of a red pellet the time was recorded, and the mice were returned to their home cages, with *ad lib* access to food and water. Any mice that did not produce a red fecal pellet were returned to their cages at 8hrs post-gavage and the time recorded was set to a maximum value of 8.5hrs for their transit time.

### Ex vivo colonic contractility recordings

Tests were performed at 2 weeks post laminectomy or spinal cord injury in independent cohorts selected prior to their injury. Mice were anesthetized via brief isoflurane inhalation followed by urethane (i.p., 2g/kg), decapitation and tissue collection. Whole colon together with cecum was removed and transferred to a dissection chamber containing ice-cold Krebs solution saturated with carbogen (95% O_2_, 5% CO_2_). Cecum and attached adipose tissue then were carefully dissected away from the colon, and the colon was transferred to a recording chamber containing carbogen saturated Krebs solution and maintained at 34°C, with a flow rate of ∼6ml/min. After 1h of incubation, a force transducer (dual force and length controller, 300C-LR, Aurora Scientific, Inc) was attached to the oral end of the distal colon with the distal end fixed to the recording chamber.

Data were acquired using AxoClamp 900A (Axon Instruments). Each tissue segment was recorded for at least 30min in 5min gap-free files. 15min of stable recordings were selected from the middle of each recording session for analysis. Using Clampfit software (Molecular Devices, RRID:SCR_011323), data files were filtered at 300Hz (Bessel 8-pole) and reduced by a factor of 100. Files were concatenated and the baseline was adjusted based on overall slope. Amplitude was calculated by subtracting the minimum value of the whole trace from the value of the peak being assessed.

Krebs solution was used in ex vivo colon motility experiments. It was composed of (in mM) 117 NaCl, 4.6 KCl, 2.5 CaCl_2_, 1.2 MgSO_4_, 1 NaHPO_4_, 11 D-glucose and 25 NaH_2_CO_3_. It was saturated with carbogen (95% O_2_, 5% CO_2_) and maintained at pH 7.4.

### Microbiome sequencing

Mice were placed into sterile 1000ml plastic cups in a biosafety cabinet. Fecal pellets were collected with sterilized forceps and placed into sterile 1.5ml plastic tubes. Tubes were immediately frozen over dry ice and placed into a −80°C freezer until they were shipped, on dry ice, for DNA extraction and 16S sequencing.

Full service 16S sequencing and computational analysis was performed via Zymo Inc (Irvine, CA) through the commercial ZymoBIOMICS® Targeted Sequencing Service pipeline (Zymo Research, Irvine, CA). Briefly, DNA was extracted using ZymoBIOMICS®-96 MagBead DNA Kit. Bacterial 16S ribosomal RNA gene targeted sequencing was performed using the Quick-16S™ NGS Library Prep Kit (Zymo Research, Irvine, CA). Vendor-designed primers for bacterial 16S amplified the V3-V4 region, pooled for equal molarity, cleaned with the Select-a-Size DNA Clean & Concentrator™ (Zymo Research, Irvine, CA), then quantified with TapeStation® (Agilent Technologies, Santa Clara, CA) and Qubit® (Thermo Fisher Scientific, Waltham, WA). The ZymoBIOMICS® Microbial Community Standard (Zymo Research, Irvine, CA) was used as a positive control for each DNA extraction and the ZymoBIOMICS® Microbial Community DNA Standard (Zymo Research, Irvine, CA) was used as a positive control for each targeted library preparation. Negative controls (*i.e.* blank extraction control, blank library preparation control) were included by the vendor to assess the level of bioburden carried by the wet-lab process. The final library was sequenced on Illumina® MiSeq™ with a V3 reagent kit (600 cycles). The sequencing was performed with 10% PhiX spike-in. Unique amplicon sequences variants were inferred from raw reads using the DADA2 pipeline^54^. Potential sequencing errors and chimeric sequences were also removed with the DADA2 pipeline. Taxonomy assignment was performed using Uclust from Qiime v.1.9.1 with the Zymo Research Database. Composition visualization, alpha-diversity, and beta-diversity analyses were performed with Qiime v.1.9.1^55^ and visualized using GraphPad PRISM software.

### SCFA analysis

*<u>Fecal samples</u>* were collected into 1.5ml plastic tubes with 500µL of methanol, over dry ice, and stored at −80°C until analysis by the Emory Lipidomics Core. Samples were homogenized in 50% acetonitrile with disruptor beads, then centrifuged at 4000xg for 10 min at 4°C. 40µL of supernatant from each sample was further derivatized with 20µL 200mM 3-Nitrophenylhydrzine, 20µL 120mM N-(3-dimethylaminopropyl)-N’-ethylcarodimmide, and 20µL 6% pyridine for 30 min at 40°C. 1.5mL of 10% acetonitrile was added to stop the reaction. The derivatized solution was then filtered and injected into LCMS for further data analysis. 10µL of each sample was injected into a mass spectrometer (Sciex 5500) to generate data. Lipid samples were passed over an Accucore C18 (4.6 x 100mm, 2.6µm) analytical column at 40°C for separation with aqueous mobile phase consisting of 0.1% formic acid in water and organic phase consisting of 0.1% formic acid in acetonitrile. The SCFA were analyzed with multiple reaction monitoring scans. A pool QC and a standard curve were run after every 10 samples to ensure the quality of the sample analysis as well as the instrument performance. Standards consisted of Acetate, Formate, Propionate, Butyrate, Valerate, Stearic, Palmitic, Oleic, Linoleic, Arachidonic, and Linolenic fatty acids. The concentration of the detected SCFA species were determined based on 6 points calibration curves using external standards with R square value greater than 0.95.

*<u>Serum samples</u>* were collected at experimental endpoint and stored at −80°C until analysis. Samples were analyzed by Metabolon, Inc for eight short chain fatty acids: acetic acid (C2), propionic acid (C3), isobutyric acid (C4), butyric acid (C4), 2-methyl-butyric acid (C5), isovaleric acid (C5), valeric acid (C5) and caproic acid (hexanoic acid, C6) by LC-MS/MS. Samples were spiked with stable labelled internal standards and subjected to protein precipitation with an organic solvent. After centrifugation, an aliquot of the supernatant was derivatized and injected into an Agilent 1290/SCIEX QTRAP 5500 LC-MS/MS system equipped with a C18 reverse-phase UHPLC column. The mass spectrometer was operated in negative mode using electrospray ionization (ESI). The peak area of the individual analyte product ions was measured against the peak area of the product ions of the corresponding internal standards. Quantitation was performed using a weighted linear least squares regression analysis generated from fortified calibration standards prepared concurrently with study samples. LC-MS/MS raw data was collected using SCIEX software Analyst 1.7.3 and processed using SCIEX OS-MQ software v3.1.6. Data reduction was performed using Microsoft Excel for Microsoft 365 MSO.

### Western blots

500µl of ice-cold Meso Scale Discovery (MSD) homogenization buffer (1L of buffer: 125ml of 1M Tris, 30ml 0.5M MgCl, 25ml of 0.1M EDTA, 10ml Triton X 100, 810ml deionized water [addition of 1 tablet of protease inhibitor (Thermo, A32961) per 10ml of buffer]) was added to frozen tissue, which was then quickly dissolved over ice using a probe sonicator. Lysed samples were then centrifuged for 10 min at 20,000x *g* at 4°C and the soluble, protein-rich supernatant was moved into a fresh 1.5ml tube. Protein levels were quantified using a standard Pierce BCA protein assay kit (Thermo Scientific). Samples were normalized to 1.5µg/µl with Novex Tris-glycine SDS sample buffer with 10% BME. Samples were denatured by boiling for 15 minutes and separated on a 15-well Novex WedgeWell 4-20% Tris-Glycine Gel (Thermo Scientific) before transfer to a 0.22µm PVDF membrane overnight at 4°C. Membranes were then blocked for 2hrs in a 5% bovine serum albumin (BSA) solution in Tris-buffered saline with 0.1% Tween-20 (TBST). Following blocking, primary antibodies diluted in blocking solution (Supplementary Table S6), were incubated on membrane overnight at 4°C. Membranes were washed in TBST and incubated at RT with indicated diluted secondary antibodies (Supplementary Table S6), conjugated to horseradish peroxidase enzyme. Chemiluminescence images were taken using an Azure Biosystems c400 imaging system and Cell Signaling SignalFire ECL reagent. Images were then quantified using FIJI, and normalized to a loading control, either GAPDH, beta actin, or total protein (quantified via Coomassie Brilliant Blue staining), as indicated in each figure.

### Multiplexed ELISAs

Tissue was prepared via sonication in MSD homogenization buffer, detailed above. 50µl of each sample, at a protein concentration of 1.6µg/µl for tissue lysate, or undiluted serum, were analyzed using the V-PLEX proinflammatory panel 1 mouse kit (MSD, K15048D-2) and the U-PLEX metabolic hormone mouse panel combo 1 (MSD, K15306K-1). Assays were performed following manufacture’s protocol and analyzed on the MSD QuickPlex SQ120 instrument and evaluated on the MSD discovery workbench 4.0 platform.

### Immunofluorescence imaging

Following tissue fixation (described above), full-thickness colon segments of ∼1cm in length were blocked for 2 hrs in 1.5ml tubes containing 1 drop of Mouse-on-Mouse blocking reagent (MOM; Vector Labs) in 1ml of PBS with 0.5% Triton-X 100. Tissue was then added to 1ml of primary antibody mixture containing primary antibodies (Supplementary Table S6) diluted in 3% normal goat serum (NGS) and 0.1% Triton X 100 in PBS, for ∼72hrs with gentle agitation at 4°C. The tissue was then washed 5 times in PBS for 1hr each with gentle agitation at room temperature. Tissue was then placed into amber 1.5ml tubes containing secondary antibodies (Supplementary Table S6) diluted in in 3% NGS and 0.1% Triton X 100 in PBS, overnight with gentle agitation at room temperature. The tissue was then washed 3 times for 1hr each, protected from light. The tissue was then stained with DAPI (1:300 in PBS) in 1.5ml tubes for 1hr at RT with gentle shaking, followed by 3 more 1hr washes in PBS. Tissue segments were then placed in 1ml of refractive index matching solution (RIMS)^56^ buffer (88% Histodenz [Sigma, D2158] in 0.02M phosphate buffer with 0.1% Tween-20 and 0.01% sodium azide) overnight at room temp with gentle agitation. Following tissue clearing, samples were mounted in 100µl of fresh RIMS, luminal side down, on glass slides with 0.25mm tissue spacers (SunJin labs). Glass cover slips were sealed to the spacers with clear nail polish.

Slides were imaged with a Leica SP8 multiphoton confocal microscope with Leica Application Suite software. Images were collected using Z-stacks of the myenteric plexus. Cells and ganglia were quantified by a blinded experimenter using FIJI Image J and the Gut Analysis Toolbox plugin^57^ for semi-automated detection of HuC/D^+^ cell bodies. Ganglia were defined as clusters of 4 or more HuC/D^+^ or PGP9.5^+^ cell bodies separated by a distance of 4 cell bodies from any other cluster. Ganglia were quantified by two separate researchers and averaged. 2-7 colonic regions were assessed per mouse, with 4-8 mice in each group.

### Bacterial manipulations and colonization

Bacteria were obtained from the American Type Culture Collection (ATCC) and cultured in the indicated conditions in Supplementary Table S6. For mono-colonization, overnight cultures were pelleted and resuspended in 50% glycerol:PBS and a single 100µl gavage provided to male and female GF mice. Colonization was confirmed via fecal culture on indicated selective media under aerobic and anaerobic conditions (5% H_2_, 10% CO_2_, 85% N_2_). Probiotic bacterial administration occurred daily. For whole microbiome reconstitution, fecal pellets were resuspended in sterile PBS with 5% sodium bicarbonate and a single 100µl gavage administered to male and female GF mice.

## Data Availability

All raw numerical data, statistical outputs, western blots are available in the Source Data files associated with this manuscript.

## Acknowledgements

We thank Rodger Liddle, Marie-Claude Perrault, and Maureen Sampson for critical reading of this manuscript and Karmarcha Martin, Shangrila Parvin, Justin Kim, Isabel Fraccaroli, Nicholas Au Yong, and all the members of the Division of Animal Resources for technical support. We acknowledge support from the Emory Gnotobiotic Animal Core (EGAC), the Emory Multiplexed Immunoassay Core (EMIC), the Emory Integrated Metabolomics & Lipidomics Core (EIMLC) and the Emory Integrated Cellular Imaging Core (ICI) which are subsidized by the Emory University School of Medicine as Integrated Core Facilities and are supported by the Georgia Clinical and Translational Science Alliance of the NIH (UL1TR002378). This work was supported by the following grants to TRS: Craig H Neilsen Foundation #642928 and NIH/NIEHS R01ES032440. AMH was supported by NIH/NINDS T32 grant NS096050-24, LBR by NIH/NIA F31AG076332, and SS by NIH/NIDDK R01DK080684. The content is solely the responsibility of the authors and does not necessarily reflect the official views of the sponsors.

## Author information

### Contributions

AMH and TRS conceptualized the study and research plan. AMH performed the animal surgeries. AMH, JC, and LBR performed the gnotobiotic experiments. YL and AMH performed and analyzed the ex vivo recordings. AMH performed and analyzed all behavioral, molecular, and microbiome assessments. SK, AMH, NK, and AW performed and analyzed intestinal histology. SS and SG provided critical expertise and conceptual discussion. AMH and TRS wrote the manuscript. All authors revised the manuscript. TRS supervised the study.

### Corresponding author

Correspondence to Timothy R. Sampson

## Supplementary Figure Legends

**Supplementary Figure S1, Related to Figure 2. Cohort-dependent impacts on SCI-triggered dysbiosis.**

16S sequencing results from an independent cohort of male mice. **A, B,** Pre-surgery to end point (14-dpi) changes in fecal microbiome 16S rRNA alpha diversity represented by Chao1(**A**) and Shannon (**B**) diversity measures. **C**, Principal component analysis (PCA) plot representing changes in overall community structure of fecal microbiomes. **D**, Microbial composition barplot, order level, of fecal microbiomes at 14-dpi. Asterisks in (**A)** and (**B)** represent a significant decrease in alpha diversity from 0-dpi to 14-dpi in the sham group. Each circle represents the average of all mice in a group for all but **C**, where each circle represents an individual mouse. N=3-8. * *P* < 0.05. Data are shown as mean ± SEM. Statistically significant differences were determined by repeated measures 2-way ANOVA with post-hoc Sidak’s multiple comparison test to compare temporal changes within groups.

**Supplementary Figure S2, Related to Figure 3. Injury and diet-associated microbes are not sufficient to trigger NBD.**

**A-C**, Heatmap of cytokines present in proximal colon tissue lysate from male and female uninjured germ-free mice that were colonized with either sham-derived or SCI-veh-derived fecal microbiomes (**A**), as assessed by multiplexed ELISA, with significant increases in CXCL1 (**B**) and IL-12p70 (**C**). **D-G**, Heatmap of serum metabolic markers, as assessed by multiplexed ELISA (**D**), with significant decreases in C-peptide (**E**), TNF (**F**), and ghrelin (**G**). **H-M**, Heatmap of serum metabolic markers in male mice with laminectomy (Sham) or injury (SCI-veh), as assessed by multiplexed ELISA (**H**), with significant decreases in C-peptide (**I**), TNF (**J**), glucagon (**K**), GLP-1 (**L**), and leptin (**M**). Each data point represents one mouse. Squares represent ex-GF males, hexagons represent ex-GF females, and circles represent SPF males. N=19-21 (**A-C**), N=14-15 (**D-G),** N=11-14 (**H-M**). * *P* < 0.05, ** *P* < 0.01, *** *P* < 0.001, **** *P* < 0.0001. Data are shown as mean ± SEM. Statistically significant differences were determined by two-tailed unpaired t-test.

**Supplementary Figure S3, Related to Figure 4. SCFA-induced IL-10 signaling is necessary for SCI recovery.**

**A**, Heatmap of cytokines, assessed by multiplexed ELISA, in proximal colon tissue lysate from male wild-type mice with laminectomy (Sham), SCI with standard diet (SCI-vehicle), and SCI with inulin-supplemented diet (SCI-inulin). **B**, Concentration of colonic IL-1β by ELISA. **C-E,** Endpoint quantification of indicated short-chain fatty-acids from fecal pellets of male IL10rb KO mice with SCI (SCI-veh) or inulin-supplemented diet (SCI-inulin). **F**, Quantification of colonic free fatty acid receptor 2 (FFAR2) levels, by western blot, in male IL10rb KO mice with SCI. **G**, Heatmap of cytokines, assessed by multiplexed ELISA, in serum from male wild type mice with SCI treated with either vehicle (SCI-veh), tripropionin (SCI-TRP), or tributyrin (SCI-TRB). **H**, Concentration of serum IL-6 by ELISA. **I-K,** Endpoint quantification of indicated short-chain fatty-acids from serum of male mice with SCI, treated with vehicle (Veh), tripropionin (TRP), or tributyrin (TRB). Each data point represents one mouse. N=11-17 (**A, B**), N=7-10 (**C-F**), N=11-13 (**G, H**), N=5-6 (**I-K**). * *P* < 0.05, ** *P* < 0.01. Data are shown as mean ± SEM. Statistically significant differences were determined by ordinary one-way ANOVA with post-hoc Tukey’s (**A, B**) or Dunnett’s (**G, K**) multiple comparison test. Or by two-tailed unpaired t-test (**C-F**).

