## Supplemental Figures 1-3 for "Diet-microbiome interactions promote enteric nervous system resilience following spinal cord injury"

**Supplementary Figure S1. Cohort-dependent impacts on SCI-triggered dysbiosis**

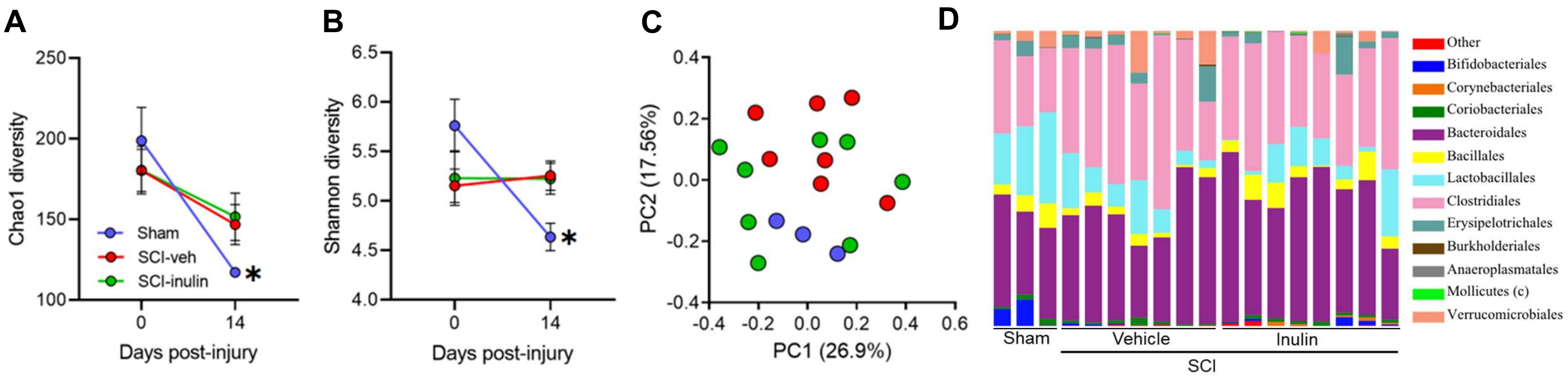

**Supplementary Figure S2. Injury and diet-associated microbes are not sufficient to trigger NBD**

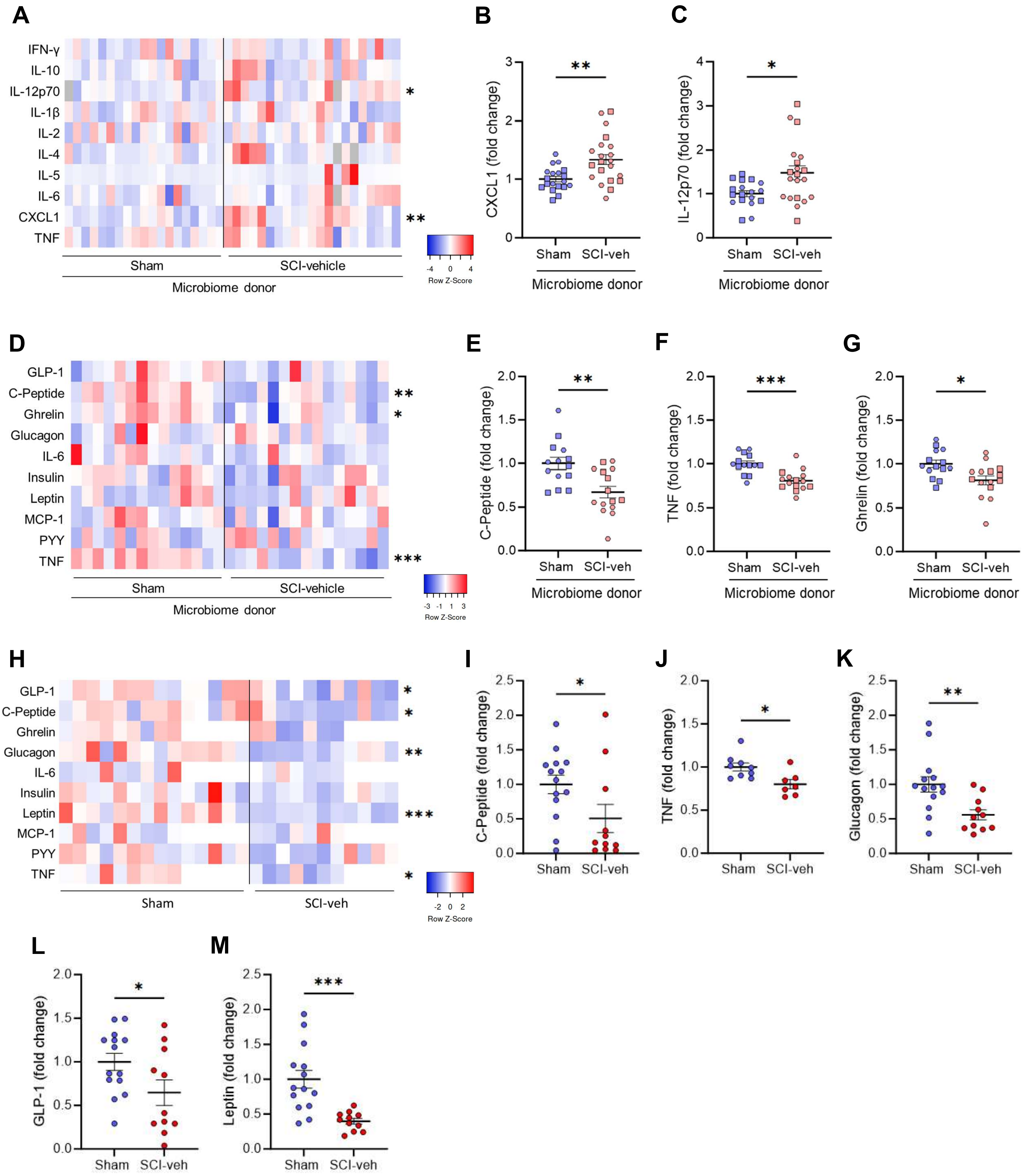

**Supplementary Figure S3. SCFA-induced IL-10 signaling is necessary for SCI recovery**

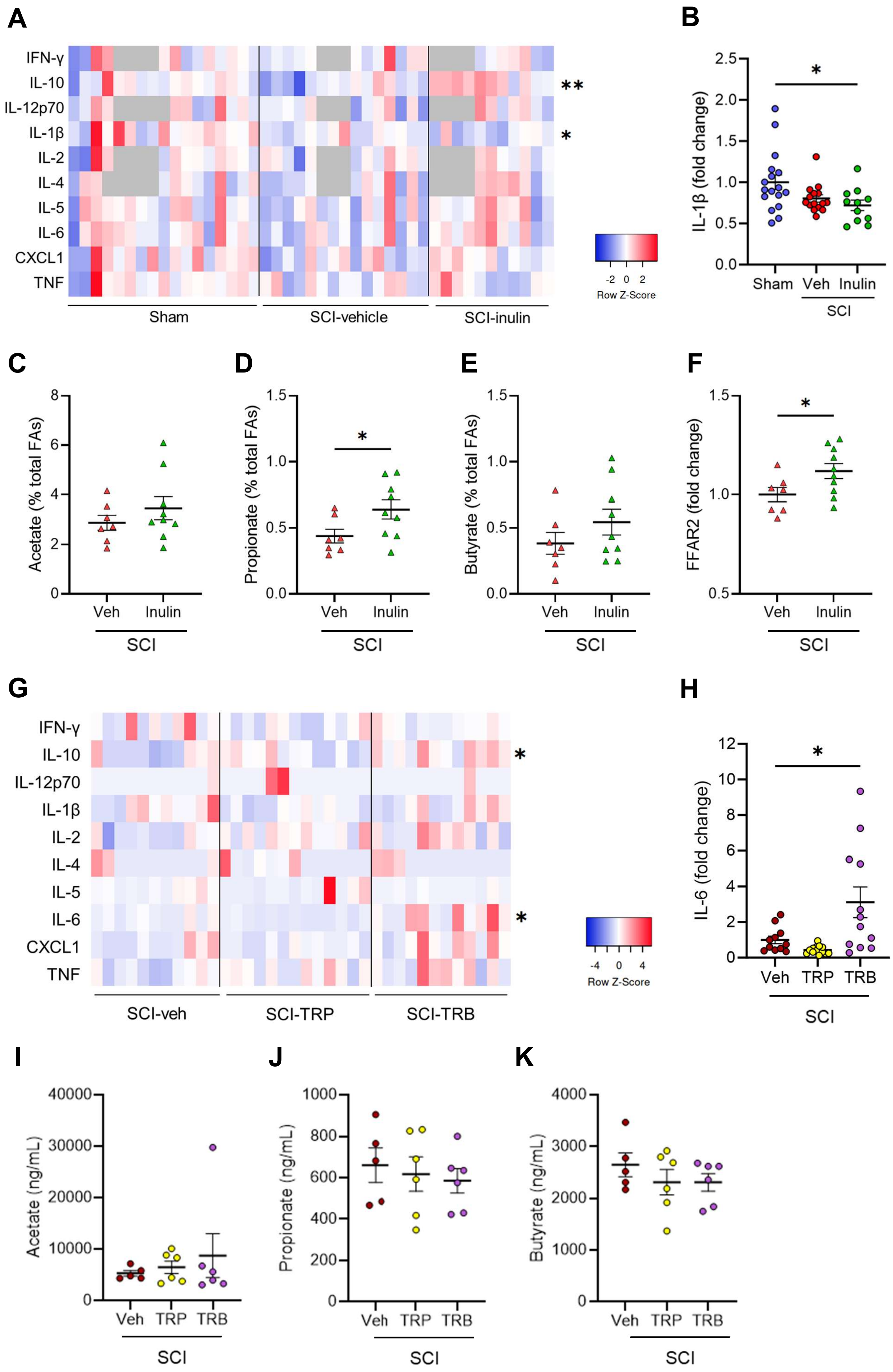
