## Supplemental Tables 1-6 for "Diet-microbiome interactions promote enteric nervous system resilience following spinal cord injury"

**Table S1. SCI-induced microbiome compositional differences.** ASV-level taxonomic changes from stool microbiomes at 2 weeks post-SCI or sham, with 1% FDR

| ***Endpoint*** | | | |
|---|---|---|---|
| **Sham vs SCI-Vehicle** | | | |
| **ASV** | **Sham** | **SCI-Veh** | **Adjusted *p* value (1% FDR)** |
| *Bacteroides sp12288* (*thetaiotaomicron*) | 0.0448409 | 0.0186638 | 6.94559E-07 |
| *Bacteroidales sp12656* | 0.156247 | 0.0943178 | 1.16422E-30 |
| *Staphylococcus lentus* | 0.0334159 | 0.00351487 | 1.50293E-08 |
| *Lactobacillus johnsonii* | 0.0577711 | 0.0290364 | 5.24441E-08 |
| *Clostridium celatum* | 0.0875609 | 0 | 0 |
| *Lachnospiraceae sp33565* | 0.00720456 | 0.0506877 | 2.82898E-16 |
| *Turicibacter sanguinis* | 0.124592 | 0.0603193 | 8.76368E-33 |
| **Sham vs SCI-Inulin** | | | |
| **ASV** | **Sham** | **SCI-Inulin** | **Adjusted *p* value (1% FDR)** |
| *Bacteroides sp12288* (*thetaiotaomicron*) | 0.0448409 | 0.0757577 | 5.81151E-08 |
| *Bacteroidales sp12550* | 0.138146 | 0.028615 | 0 |
| *Bacteroidales sp12656* | 0.156247 | 0.188446 | 1.62892E-08 |
| *Staphylococcus lentus* | 0.0334159 | 0.000353782 | 6.74178E-09 |
| *Clostridium celatum* | 0.0875609 | 0 | 0 |
| *Lachnospiraceae sp33565* | 0.00720456 | 0.040268 | 6.73239E-09 |
| *Turicibacter sanguinis* | 0.124592 | 0 | 0 |
| **SCI-Vehicle vs SCI-Inulin** | | | |
| **ASV** | **SCI-Veh** | **SCI-Inulin** | **Adjusted *p* value (1% FDR)** |
| *Bacteroides sp12288* (*thetaiotaomicron*) | 0.0186638 | 0.0757577 | 1.15921E-15 |
| *Bacteroidales sp12550* | 0.135412 | 0.028615 | 0 |
| *Bacteroidales sp12656* | 0.0943178 | 0.188446 | 2.24226E-38 |
| *Lactobacillus johnsonii* | 0.0290364 | 0.061178 | 5.5047E-06 |
| *Turicibacter sanguinis* | 0.0603193 | 0 | 2.91586E-17 |
| **Pre-SCI vs Post-SCI** | | | |
| **ASV** | **SCI-Pre** | **SCI-2wks** | **Adjusted *p* value (1% FDR)** |
| *Bacteroidales sp12656* | 0.158599 | 0.0943178 | 2.06832E-30 |
| *Lactobacillus johnsonii* | 0.193283 | 0.0290364 | 0 |
| *Clostridium celatum* | 0.022866 | 0 | 2.92759E-05 |
| *Lachnospiraceae sp33565* | 0.0128364 | 0.0506877 | 6.26693E-12 |
| *Turicibacter sanguinis* | 0.00972012 | 0.0603193 | 7.71416E-20 |

**Table S2. LEfSE identified taxonomic changes from stool microbiomes at 2 weeks post-SCI or sham**

| Bacterial taxa | Group | Effect size | *p*-value |
|---|---|---|---|
| k__Bacteria.p__Firmicutes.c__Clostridia.o__Clostridiales.f__Lachnospiraceae.g__NA.s__sp33760 | SCI.Inulin | 3.28014007 | 0.030197383 |
| k__Bacteria.p__Firmicutes.c__Clostridia.o__Clostridiales.f__NA.g__NA.s__sp31072 | SCI.Inulin | 2.98922362 | 0.048484305 |
| k__Bacteria.p__Proteobacteria.c__Gammaproteobacteria.o__Pseudomonadales.f__Moraxellaceae.g__Acinetobacter.s__radioresistens | SCI.Inulin | 3.18049855 | 0.012764197 |
| k__Bacteria.p__Firmicutes.c__Clostridia.o__Clostridiales.f__Lachnospiraceae.g__NA.s__sp32706 | SCI.Inulin | 3.82883697 | 0.01334429 |
| k__Bacteria.p__Firmicutes.c__Clostridia.o__Clostridiales.f__Lachnospiraceae.g__NA.s__sp32638 | SCI.Inulin | 3.78461598 | 0.023383895 |
| k__Bacteria.p__Firmicutes.c__Clostridia.o__Clostridiales.f__Lachnospiraceae.g__NA.s__sp33456 | SCI.Inulin | 3.17262743 | 0.027158625 |
| k__Bacteria.p__Firmicutes.c__Clostridia.o__Clostridiales.f__Lachnospiraceae.g__NA.s__sp33694 | SCI.Inulin | 3.76036576 | 0.019757099 |
| k__Bacteria.p__Firmicutes.c__Clostridia.o__Clostridiales.f__Ruminococcaceae.g__NA.s__sp34867 | SCI.Inulin | 3.30028204 | 0.027323722 |
| k__Bacteria.p__Firmicutes.c__Clostridia.o__Clostridiales.f__Ruminococcaceae.g__Anaerotruncus.s__sp34475 | SCI.Inulin | 2.99522399 | 0.045806314 |
| k__Bacteria.p__Proteobacteria.c__Gammaproteobacteria.o__Pseudomonadales | SCI.Inulin | 3.25550068 | 0.016389554 |
| k__Bacteria.p__Proteobacteria.c__Gammaproteobacteria.o__Pseudomonadales.f__Moraxellaceae.g__Acinetobacter | SCI.Inulin | 3.22914314 | 0.016389554 |
| k__Bacteria.p__Firmicutes.c__Clostridia.o__Clostridiales.f__Lachnospiraceae.g__NA.s__sp33417 | SCI.Inulin | 3.75893236 | 0.011625603 |
| k__Bacteria.p__Bacteroidetes.c__Bacteroidia.o__Bacteroidales.f__Bacteroidaceae.g__Bacteroides.s__sp12288_thetaiotaomicron | SCI.Inulin | 4.43785167 | 0.028207093 |
| k__Bacteria.p__Firmicutes.c__Clostridia.o__Clostridiales.f__Lachnospiraceae.g__NA.s__sp33525 | SCI.Inulin | 3.26973262 | 0.048484305 |
| k__Bacteria.p__Firmicutes.c__Clostridia.o__Clostridiales.f__Lachnospiraceae.g__NA.s__sp32701 | SCI.Inulin | 3.96461728 | 0.015562692 |
| k__Bacteria.p__Firmicutes.c__Clostridia.o__Clostridiales.f__Lachnospiraceae.g__NA.s__sp33616_sp33639 | SCI.Inulin | 3.42818688 | 0.029407217 |
| k__Bacteria.p__Firmicutes.c__Bacilli.o__Lactobacillales.f__Enterococcaceae.g__Enterococcus.s__faecalis | SCI.Inulin | 3.05716304 | 0.024861262 |
| k__Bacteria.p__Proteobacteria.c__Gammaproteobacteria.o__Pseudomonadales.f__Moraxellaceae.g__Acinetobacter.s__calcoaceticus_radioresistens_variabilis | SCI.Inulin | 3.20043929 | 0.041388896 |
| k__Bacteria.p__Firmicutes.c__Clostridia.o__Clostridiales.f__Ruminococcaceae.g__NA.s__sp34731 | SCI.Inulin | 3.43239165 | 0.027158625 |
| k__Bacteria.p__Firmicutes.c__Bacilli.o__Lactobacillales.f__Enterococcaceae | SCI.Inulin | 3.0725174 | 0.024861262 |
| k__Bacteria.p__Bacteroidetes.c__Bacteroidia.o__Bacteroidales.f__Bacteroidaceae | SCI.Inulin | 4.41723887 | 0.028207093 |
| k__Bacteria.p__Bacteroidetes.c__Bacteroidia.o__Bacteroidales.f__Bacteroidaceae.g__Bacteroides | SCI.Inulin | 4.45069433 | 0.028207093 |
| k__Bacteria.p__Firmicutes.c__Clostridia.o__Clostridiales.f__Lachnospiraceae.g__NA.s__sp32758 | SCI.Inulin | 3.97272763 | 0.009403563 |
| k__Bacteria.p__Firmicutes.c__Clostridia.o__Clostridiales.f__Lachnospiraceae.g__NA.s__sp32623 | SCI.Inulin | 3.56178608 | 0.041041078 |
| k__Bacteria.p__Firmicutes.c__Clostridia.o__Clostridiales.f__NA.g__NA.s__sp31116 | SCI.Inulin | 3.05573343 | 0.04376775 |
| k__Bacteria.p__Proteobacteria.c__Gammaproteobacteria.o__Pseudomonadales.f__Moraxellaceae | SCI.Inulin | 3.23322001 | 0.016389554 |
| k__Bacteria.p__Firmicutes.c__Bacilli.o__Lactobacillales.f__Enterococcaceae.g__Enterococcus | SCI.Inulin | 3.06357795 | 0.024861262 |
| k__Bacteria.p__Firmicutes.c__Clostridia.o__Clostridiales.f__Lachnospiraceae.g__NA.s__sp33721 | SCI.Inulin | 3.08898072 | 0.009139753 |
| k__Bacteria.p__Firmicutes.c__Clostridia.o__Clostridiales.f__Ruminococcaceae.g__NA.s__sp35841 | SCI | 3.08050444 | 0.030197383 |
| k__Bacteria.p__Firmicutes.c__Clostridia.o__Clostridiales.f__Lachnospiraceae.g__NA.s__sp33658 | SCI | 3.42184273 | 0.023786517 |
| k__Bacteria.p__Firmicutes.c__Clostridia.o__Clostridiales.f__Lachnospiraceae.g__NA.s__sp32702 | SCI | 3.85317688 | 0.024861262 |
| k__Bacteria.p__Firmicutes.c__Clostridia.o__Clostridiales.f__Lachnospiraceae.g__Lachnoclostridium.s__sp32402 | SCI | 3.69380202 | 0.034609178 |
| k__Bacteria.p__Firmicutes.c__Clostridia.o__Clostridiales.f__Lachnospiraceae.g__NA.s__sp33739 | SCI | 3.15551336 | 0.046329809 |
| k__Bacteria.p__Firmicutes.c__Clostridia.o__Clostridiales.f__Lachnospiraceae | SCI | 5.05697495 | 0.048668305 |
| k__Bacteria.p__Firmicutes.c__Clostridia.o__Clostridiales.f__Lachnospiraceae.g__NA.s__sp33565 | SCI | 4.28952463 | 0.027158625 |
| k__Bacteria.p__Firmicutes.c__Clostridia.o__Clostridiales.f__Lachnospiraceae.g__NA.s__sp32872 | SCI | 3.09241069 | 0.028207093 |
| k__Bacteria.p__Firmicutes.c__Clostridia.o__Clostridiales.f__Ruminococcaceae.g__Oscillibacter.s__sp34648 | SCI | 3.04716797 | 0.028937471 |
| k__Bacteria.p__Firmicutes.c__Clostridia.o__Clostridiales.f__Lachnospiraceae.g__NA.s__sp33718 | SCI | 3.35285573 | 0.023786517 |
| k__Bacteria.p__Firmicutes.c__Clostridia.o__Clostridiales.f__Lachnospiraceae.g__NA.s__sp32146 | SCI | 3.27855544 | 0.027158625 |
| k__Bacteria.p__Firmicutes.c__Clostridia.o__Clostridiales.f__Ruminococcaceae.g__NA.s__sp35077 | Sham | 3.08269556 | 0.027323722 |
| k__Bacteria.p__Firmicutes.c__Bacilli.o__Bacillales.f__Staphylococcaceae.g__Staphylococcus.s__lentus | Sham | 4.22428258 | 0.015562692 |

**Table S2 (continued)**

| Bacterial taxa | Group | Effect size | *p*-value |
|---|---|---|---|
| k__Bacteria.p__Firmicutes.c__Erysipelotrichia.o__Erysipelotrichales | Sham | 4.77308354 | 0.015562692 |
| k__Bacteria.p__Firmicutes.c__Clostridia.o__Clostridiales.f__Clostridiaceae | Sham | 4.65015938 | 0.025161049 |
| k__Bacteria.p__Firmicutes.c__Clostridia.o__Clostridiales.f__Clostridiaceae.g__Clostridium.s__celatum | Sham | 4.6301129 | 0.009139753 |
| k__Bacteria.p__Firmicutes.c__Clostridia.o__Clostridiales.f__Ruminococcaceae.g__NA.s__sp35361 | Sham | 3.59813998 | 0.019660049 |
| k__Bacteria.p__Firmicutes.c__Clostridia.o__Clostridiales.f__Ruminococcaceae.g__NA.s__sp35360 | Sham | 3.08921088 | 0.041041078 |
| k__Bacteria.p__Proteobacteria.c__Betaproteobacteria.o__Burkholderiales.f__Alcaligenaceae | Sham | 3.43800153 | 0.011391522 |
| k__Bacteria.p__Firmicutes.c__Bacilli.o__Bacillales.f__Staphylococcaceae.g__Staphylococcus | Sham | 4.27885163 | 0.037902914 |
| k__Bacteria.p__Firmicutes.c__Clostridia.o__Clostridiales.f__Lachnospiraceae.g__NA.s__sp33419 | Sham | 3.32307121 | 0.015562692 |
| k__Bacteria.p__Bacteroidetes.c__Bacteroidia.o__Bacteroidales.f__NA.g__NA.s__sp12550 | Sham | 4.72904022 | 0.048484305 |
| k__Bacteria.p__Firmicutes.c__Clostridia.o__Clostridiales.f__Clostridiaceae.g__Clostridium | Sham | 4.65240631 | 0.025161049 |
| k__Bacteria.p__Firmicutes.c__Erysipelotrichia.o__Erysipelotrichales.f__Erysipelotrichaceae.g__Turicibacter.s__sanguinis | Sham | 4.7786212 | 0.012764197 |
| k__Bacteria.p__Firmicutes.c__Bacilli.o__Bacillales.f__Staphylococcaceae.g__Jeotgalicoccus | Sham | 3.42659395 | 0.013728404 |
| k__Bacteria.p__Firmicutes.c__Bacilli.o__Bacillales.f__Staphylococcaceae | Sham | 4.30010345 | 0.037902914 |
| k__Bacteria.p__Firmicutes.c__Erysipelotrichia | Sham | 4.76829196 | 0.015562692 |
| k__Bacteria.p__Firmicutes.c__Clostridia.o__Clostridiales.f__Lachnospiraceae.g__NA.s__sp33537 | Sham | 3.08288654 | 0.041388896 |
| k__Bacteria.p__Firmicutes.c__Clostridia.o__Clostridiales.f__NA.g__NA.s__sp31125 | Sham | 3.07776853 | 0.041388896 |
| k__Bacteria.p__Firmicutes.c__Clostridia.o__Clostridiales.f__NA.g__NA.s__sp31122 | Sham | 3.42566575 | 0.034411331 |
| k__Bacteria.p__Firmicutes.c__Erysipelotrichia.o__Erysipelotrichales.f__Erysipelotrichaceae.g__Turicibacter | Sham | 4.801587 | 0.012764197 |
| k__Bacteria.p__Firmicutes.c__Erysipelotrichia.o__Erysipelotrichales.f__Erysipelotrichaceae | Sham | 4.77876752 | 0.015562692 |
| k__Bacteria.p__Firmicutes.c__Clostridia.o__Clostridiales.f__Ruminococcaceae.g__NA.s__sp35498 | Sham | 3.08729405 | 0.041388896 |
| k__Bacteria.p__Proteobacteria.c__Betaproteobacteria.o__Burkholderiales | Sham | 3.41658553 | 0.011391522 |
| k__Bacteria.p__Proteobacteria.c__Betaproteobacteria | Sham | 3.40401102 | 0.011391522 |
| k__Bacteria.p__Firmicutes.c__Bacilli.o__Bacillales | Sham | 4.30946747 | 0.037902914 |
| k__Bacteria.p__Proteobacteria.c__Betaproteobacteria.o__Burkholderiales.f__Alcaligenaceae.g__Parasutterella | Sham | 3.41352673 | 0.011391522 |
| k__Bacteria.p__Firmicutes.c__Bacilli.o__Bacillales.f__Staphylococcaceae.g__Jeotgalicoccus.s__halophilus_halotolerans_nanhaiensis | Sham | 3.4063761 | 0.013728404 |
| k__Bacteria.p__Proteobacteria.c__Betaproteobacteria.o__Burkholderiales.f__Alcaligenaceae.g__Parasutterella.s__excrementihominis | Sham | 3.42627509 | 0.011391522 |

**Table S3. Bacterial taxa significantly different following SCI in an independent cohort.** With 1% FDR correction, alterations between Sham and SCI (a), sham and SCI-inulin (b), or SCI and SCI-inulin (c). Grey boxes indicate significant comparisons (1% FDR) between pre and post-SCI groups at 2wks post injury.

| ASV | Sham | SCI-Veh | SCI-Inulin | Sig comparisons by 1% FDR | Identified in independent cohort (Fig 2) |
|---|---|---|---|---|---|
| *Akkermansia muciniphila* | 0.032851 | 0.04787 | 0.016194 | c | |
| *Alistipes sp14336* | 0.029102 | 0.067027 | 0.06584 | a, b | |
| *Allobaculum sp36555* | 0.000152 | 0.021406 | 0.006325 | a | |
| *Bacteroidales sp12473-sp12526-sp12633* | 0.074217 | 0.084304 | 0.042379 | b, c | |
| *Bacteroidales sp12610* | 0 | 0.030116 | 0.013937 | a | |
| *Bacteroidales sp12645* | 0 | 0.019732 | 0.058406 | b, c | |
| *Bacteroidales sp12656* | 0.105666 | 0.052678 | 0.156417 | a, b, c | Yes |
| *Bacteroidales sp12768* | 0.033087 | 0.05611 | 0.04598 | a | |
| *Bacteroides sp12288 (thetaiotaomicron)* | 0.080995 | 0.071137 | 0.050251 | b, c | Yes |
| *Bifidobacterium choerinum-pseudolongum* | 0.049002 | 0.002335 | 0.007556 | a, b | |
| *Lachnoclostridium sp32402* | 0.021026 | 0.040619 | 0.01123 | c | |
| *Lachnospiraceae sp32778* | 0.009123 | 0.043795 | 0.014431 | a, c | |
| *Lactobacillus johnsonii* | 0.237838 | 0.079619 | 0.078155 | a, b | Yes |
| *Staphylococcus lentus* | 0.027879 | 0.005809 | 0.019373 | a | Yes |

**Table S4. Independent SCI-induced microbiome compositional differences.** ASV-level taxonomic changes from stool microbiomes at 2 weeks post-SCI or sham, with 1% FDR, from independent cohort

| ***Endpoint*** | | | |
|---|---|---|---|
| **Sham vs SCI-Vehicle** | | | |
| **ASV** | **Sham** | **SCI-Veh** | **Adjusted *p* value (1% FDR)** |
| *Bifidobacterium choerinum-pseudolongum* | 0.0490019 | 0.00233502 | 1.38505E-15 |
| *Bacteroidales sp12610* | 0 | 0.0301156 | 2.27912E-07 |
| *Bacteroidales sp12656* | 0.105666 | 0.0526781 | 1.42237E-19 |
| *Bacteroidales sp12768* | 0.0330871 | 0.05611 | 7.46328E-05 |
| *Alistipes sp14336* | 0.0291018 | 0.0670273 | 7.71846E-11 |
| *Staphylococcus lentus* | 0.0278791 | 0.00580894 | 0.000146098 |
| *Lactobacillus johnsonii* | 0.237838 | 0.0796191 | 0 |
| *Lachnospiraceae sp32778* | 0.00912332 | 0.0437946 | 2.64513E-09 |
| *Allobaculum p36555* | 0.00015199 | 0.0214064 | 0.000254523 |
| **Sham vs SCI-Inulin** | | | |
| **ASV** | **Sham** | **SCI-Inulin** | **Adjusted *p* value (1% FDR)** |
| *Bifidobacterium choerinum-pseudolongum* | 0.0490019 | 0.00755608 | 4.74124E-08 |
| *Bacteroides sp12288 (thetaiotaomicron)* | 0.0809949 | 0.0502512 | 4.99359E-05 |
| *Bacteroidales sp12473-sp12526-sp12633* | 0.0742173 | 0.0423789 | 2.67067E-05 |
| *Bacteroidales sp12645* | 0 | 0.0584062 | 1.67097E-14 |
| *Bacteroidales sp12656* | 0.105666 | 0.156417 | 2.43458E-11 |
| *Alistipes sp14336* | 0.0291018 | 0.0658396 | 1.27625E-06 |
| *Lactobacillus johnsonii* | 0.237838 | 0.0781547 | 0 |
| **SCI-Vehicle vs SCI-Inulin** | | | |
| **ASV** | **SCI-Veh** | **SCI-Inulin** | **Adjusted *p* value (1% FDR)** |
| *Bacteroides sp12288 (thetaiotaomicron)* | 0.0711372 | 0.0502512 | 0.00013929 |
| *Bacteroidales sp12473-sp12526-sp12633* | 0.0843043 | 0.0423789 | 2.45024E-14 |
| *Bacteroidales sp12645* | 0.0197316 | 0.0584062 | 1.95902E-12 |
| *Bacteroidales sp12656* | 0.0526781 | 0.156417 | 0 |
| *Lachnoclostridium sp32402* | 0.040619 | 0.0112297 | 8.5429E-08 |
| *Lachnospiraceae sp32778* | 0.0437946 | 0.0144314 | 8.76893E-08 |
| *Akkermansia muciniphila* | 0.0478695 | 0.0161938 | 7.92208E-09 |
| **Pre SCI vs Post SCI** | | | |
| **ASV** | **SCI-Pre** | **SCI-2wks** | **Adjusted *p* value (1% FDR)** |
| *Other* | 0.0400304 | 0.00230997 | 3.01332E-20 |
| *Bifidobacterium choerinum-pseudolongum* | 0.0387627 | 0.00233502 | 5.35722E-19 |
| *Bacteroidales sp12473-sp12526-sp12633* | 0.0566063 | 0.0843043 | 1.20298E-11 |
| *Bacteroidales sp12768* | 0.0328508 | 0.05611 | 1.223E-08 |
| *Enterococcus faecalis* | 0.00133197 | 0.0180171 | 4.33168E-05 |
| *Lactobacillus johnsonii* | 0.130668 | 0.0796191 | 1.57727E-35 |
| *Lachnoclostridium sp32402* | 0.00983318 | 0.040619 | 4.98182E-14 |
| *Lachnospiraceae sp32778* | 0.00608032 | 0.0437946 | 3.05544E-20 |
| *Ruminococcaceae sp35297* | 0.0262177 | 0.0028927 | 1.11286E-08 |
| *Allobaculum sp36555* | 0.0653365 | 0.0214064 | 7.94526E-27 |

**Table S5. LEfSE identified taxonomic changes from stool microbiomes at 2 weeks post-SCI or sham, from independent cohort**

| Bacterial taxa | Group | Effect size | *p*-value |
|---|---|---|---|
| k__Bacteria.p__Firmicutes.c__Clostridia.o__Clostridiales.f__Lachnospiraceae.g__NA.s__sp33761 | SCI | 3.20746086 | 0.03938728 |
| k__Bacteria.p__Firmicutes.c__Clostridia.o__Clostridiales.f__Ruminococcaceae.g__Anaerotruncus.s__sp34475 | SCI | 3.14434302 | 0.01455464 |
| k__Bacteria.p__Bacteroidetes.c__Bacteroidia.o__Bacteroidales.f__NA.g__NA.s__sp12473_sp12526_sp12633 | SCI | 4.29010458 | 0.03319739 |
| k__Bacteria.p__Firmicutes.c__Clostridia.o__Clostridiales.f__Lachnospiraceae.g__NA.s__sp32862 | SCI | 3.49736231 | 0.03339209 |
| k__Bacteria.p__Firmicutes.c__Clostridia.o__Clostridiales.f__Lachnospiraceae.g__NA.s__sp32735 | SCI | 3.60307215 | 0.02671248 |
| k__Bacteria.p__Firmicutes.c__Clostridia.o__Clostridiales.f__Lachnospiraceae.g__NA.s__sp32721 | SCI | 3.47470577 | 0.02675007 |
| k__Bacteria.p__Firmicutes.c__Clostridia.o__Clostridiales.f__Ruminococcaceae.g__NA.s__sp33140 | SCI | 2.96559706 | 0.02508897 |
| k__Bacteria.p__Actinobacteria.c__Actinobacteria.o__Corynebacteriales.f__Corynebacteriaceae | SCI.Inulin | 3.5183703 | 0.01922687 |
| k__Bacteria.p__Firmicutes.c__Clostridia.o__Clostridiales.f__Lachnospiraceae.g__NA.s__sp33731 | SCI.Inulin | 3.21037177 | 0.01440632 |
| k__Bacteria.p__Actinobacteria.c__Actinobacteria.o__Corynebacteriales.f__Corynebacteriaceae.g__Corynebacterium | SCI.Inulin | 3.51837339 | 0.01922687 |
| k__Bacteria.p__Actinobacteria.c__Actinobacteria.o__Corynebacteriales | SCI.Inulin | 3.51837267 | 0.01922687 |
| k__Bacteria.p__Firmicutes.c__Clostridia.o__Clostridiales.f__Lachnospiraceae.g__NA.s__sp32635_sp32668 | SCI.Inulin | 3.09090389 | 0.01922687 |
| k__Bacteria.p__Actinobacteria.c__Actinobacteria.o__Corynebacteriales.f__Corynebacteriaceae.g__Corynebacterium.s__amycolatum | SCI.Inulin | 3.51837192 | 0.01922687 |
| k__Bacteria.p__Firmicutes.c__Clostridia.o__Clostridiales.f__Lachnospiraceae.g__NA.s__sp33432 | SCI.Inulin | 3.16044082 | 0.01835448 |
| k__Bacteria.p__Firmicutes.c__Clostridia.o__Clostridiales.f__Lachnospiraceae.g__Acetatifactor | SCI.Inulin | 3.0869611 | 0.0469165 |
| k__Bacteria.p__Firmicutes.c__Clostridia.o__Clostridiales.f__Lachnospiraceae.g__NA.s__sp33421_sp33679 | SCI.Inulin | 3.69414989 | 0.00122429 |
| k__Bacteria.p__Firmicutes.c__Bacilli.o__Bacillales.f__Planococcaceae.g__Sporosarcina.s__luteola_pasteurii | Sham | 3.15351433 | 0.04203087 |
| k__Bacteria.p__Firmicutes.c__Bacilli.o__Lactobacillales.f__Lactobacillaceae.g__Lactobacillus | Sham | 4.94539917 | 0.04846866 |
| k__Bacteria.p__Firmicutes.c__Bacilli.o__Lactobacillales.f__Lactobacillaceae | Sham | 4.93387968 | 0.04846866 |

**Table S5 (continued)**

| Bacterial taxa | Group | Effect size | *p*-value |
|---|---|---|---|
| k__Bacteria.p__Firmicutes.c__Bacilli.o__Bacillales.f__Staphylococcaceae.g__Staphylococcus.s__lentus | Sham | 4.05241164 | 0.02345153 |
| k__Bacteria.p__Firmicutes.c__Bacilli.o__Bacillales.f__Planococcaceae | Sham | 3.13536277 | 0.04203087 |
| k__Bacteria.p__Firmicutes.c__Erysipelotrichia.o__Erysipelotrichales.f__Erysipelotrichaceae.g__NA | Sham | 3.04568574 | 0.00943301 |
| k__Bacteria.p__Firmicutes.c__Erysipelotrichia.o__Erysipelotrichales.f__Erysipelotrichaceae.g__NA.s__sp36786 | Sham | 3.28495448 | 0.00107094 |
| k__Bacteria.p__Firmicutes.c__Clostridia.o__Clostridiales.f__Lachnospiraceae.g__NA.s__sp33694 | Sham | 3.51794408 | 0.01961457 |
| k__Bacteria.p__Firmicutes.c__Clostridia.o__Clostridiales.f__Lachnospiraceae.g__NA.s__sp33673 | Sham | 3.20052991 | 0.02425508 |
| k__Bacteria.p__Firmicutes.c__Bacilli.o__Bacillales.f__Staphylococcaceae.g__Staphylococcus | Sham | 4.23216416 | 0.04462127 |
| k__Bacteria.p__Actinobacteria | Sham | 4.407117 | 0.02774235 |
| k__Bacteria.p__Firmicutes.c__Clostridia.o__Clostridiales.f__Lachnospiraceae.g__NA.s__sp33636 | Sham | 3.49964269 | 0.00831405 |
| k__Bacteria.p__Firmicutes.c__Clostridia.o__Clostridiales.f__Lachnospiraceae.g__Marvinbryantia.s__sp32979 | Sham | 3.51749967 | 0.00322499 |
| k__Bacteria.p__Firmicutes.c__Bacilli.o__Bacillales.f__Staphylococcaceae | Sham | 4.22071125 | 0.04462127 |
| k__Bacteria.p__Firmicutes.c__Clostridia.o__Clostridiales.f__Lachnospiraceae.g__Blautia.s__sp32038 | Sham | 3.5580099 | 0.02597101 |
| k__Bacteria.p__Firmicutes.c__Clostridia.o__Clostridiales.f__Lachnospiraceae.g__NA.s__sp33374 | Sham | 3.35338706 | 0.02708786 |
| k__Bacteria.p__Firmicutes.c__Clostridia.o__Clostridiales.f__NA.g__NA.s__sp31125 | Sham | 3.0240336 | 0.00502104 |
| k__Bacteria.p__Firmicutes.c__Bacilli.o__Bacillales.f__Planococcaceae.g__Sporosarcina | Sham | 3.14293813 | 0.04203087 |
| k__Bacteria.p__Firmicutes.c__Clostridia.o__Clostridiales.f__Lachnospiraceae.g__NA.s__sp32622 | Sham | 3.84168757 | 0.03785401 |
| k__Bacteria.p__Firmicutes.c__Clostridia.o__Clostridiales.f__Lachnospiraceae.g__NA.s__sp32156 | Sham | 3.13422797 | 0.03942504 |
| k__Bacteria.p__Firmicutes.c__Clostridia.o__Clostridiales.f__Lachnospiraceae.g__Marvinbryantia | Sham | 3.63670981 | 0.0093402 |

**Supplemental Data Table 6. Reagents & Resources.**

**Antibodies used**

| Primary antibodies for Western Blot | | | | |
|---|---|---|---|---|
| **Antibody** | **Concentration** | **Host** | **Supplier** | **Cat Number** |
| Neuronal nitric oxide synthase (nNOS) | 1:1000 | Rabbit | Cell Signaling | 4231S |
| Choline acetyltransferase (ChAT) | 1:1000 | Goat | Sigma | AB144P |
| Protein gene product 9.5 (PGP9.5) | 1:1000 | Rabbit | Millipore | AB1761-I |
| Free fatty acid receptor 2 (FFAR2 / GPR43) | 1:1000 | Rabbit | Thermo | PA5-111780 |
| Glyceraldehyde 3-phosphate dehydrogenase (GAPDH) | 1:1000 | Rabbit | Cell Signaling | 5174S |
| β-Actin | 1:1000 | Rabbit | Cell Signaling | 8457S |
| **Secondary antibodies for Western Blot** | | | | |
| **Antibody** | **Concentration** | **Host** | **Supplier** | **Cat Number** |
| Anti-rabbit (HRP-linked) | 1:1000 | Goat | Cell Signaling | 7074S |
| Anti-goat (HRP-linked) | 1:1000 | Donkey | Thermo | A16005 |
| **Primary antibodies for Immunohistochemistry** | | | | |
| **Antibody** | **Concentration** | **Host** | **Supplier** | **Cat Number** |
| Neuronal nitric oxide synthase (nNOS) | 1:100 | Rabbit | Cell Signaling | 4231S |
| Anti-HuD + HuC antibody | 1:500 | Rabbit | Abcam | AB184267 |
| Protein gene product 9.5 (PGP9.5) | 1:200 | Mouse | Abcam | AB72911 |
| **Secondary antibodies for Immunohistochemistry** | | | | |
| **Antibody** | **Concentration** | **Host** | **Supplier** | **Cat Number** |
| Anti-mouse (Alexa Fluor™ 594) | 1:200 | Goat | Thermo | A-11005 |
| Anti-rabbit (Alexa Fluor™ 488) | 1:200 | Goat | Thermo | A-11008 |

**ATCC Strains and Growth Conditions**

| Bacterium | Medium | Environment | CFU per gavage | ATCC Number |
|---|---|---|---|---|
| *Lactobacillus johnsonii* | de Man-Rogosa-Sharpe | Aerobic | 10^16 | 33200 |
| *Bacteroides thetaiotaomicron* | Brain-Heart Infusion | Anaerobic | 10^14 | 29148 |
| *Clostridium celatum* | Chopped Meat Carbohydrate | Anaerobic | 10^10 | 27791 |
